# END nucleases: Antiphage defense systems targeting multiple hypermodified phage genomes

**DOI:** 10.1101/2025.03.31.646159

**Authors:** Wearn-Xin Yee, Yan-Jiun Lee, Timothy A. Klein, Alex Wirganowicz, Andres E. Gabagat, Bálint Csörgő, Kira S. Makarova, Eugene V. Koonin, Peter R. Weigele, Joseph Bondy-Denomy

## Abstract

Prokaryotes carry clusters of phage defense systems in “defense islands” that have been extensively exploited bioinformatically and experimentally for discovery of immune functions. However, little effort has been dedicated to determining which specific system(s) within defense islands limit lytic phage reproduction in clinical bacterial strains. Here, we employed the CRISPR-based Cascade-Cas3 system to delete defense islands in a *Pseudomonas aeruginosa* clinical isolate to identify mechanisms of lytic phage antagonism. Deletion of one island in a cystic fibrosis-derived clinical isolate sensitized the strain to phages from the *Pbunavirus* family, which are commonly used as therapeutics. The causal defense system is a Type IIS restriction endonuclease-like protein (END^PaCF1^), common in Pseudomonads, however it lacks an associated methyltransferase typical Type IIS R-M systems. END^PaCF1^ protects bacteria against phages with hypermodified DNA and is surprisingly agnostic to the specific structure of the modification, which is unlike typical type IV restriction endonucleases. In END^PaCF1^, the endonuclease domain is fused to a catalytically inactive Endonuclease III (iEndoIII), a domain that recognizes non-canonical bases to repair DNA in prokaryotes and eukaryotes. We therefore propose that nucleases containing an i**<u>En</u>**doIII **<u>d</u>**omain (**<u>END</u>** nucleases) can sense diverse DNA hypermodifications. Our findings reveal modularity of the sensing and cleavage domains, as expected of a modification-dependent endonucleases. We further show that some hypermodified phages, including *Pbunavirus* family members and *Wrowclawvirus* family (Pa5oct-like) of jumbo phages, encode END nuclease inhibitors that directly bind to the nuclease, likely via the iEndoIII domain. These inhibitors are necessary for *Pbunavirus* to plaque on clinical isolates and sufficient to enable other hypermodified phages to plaque in the presence of this defense system.

## Introduction

The recent discovery of numerous antiphage defense systems has opened up new research directions including the identification of a variety of novel nucleases^1^, signaling and pattern-recognition based defenses^2–4^, multiple programmed cell death mechanisms and more^5,6^. Many of these defense systems show direct evolutionary connections and/or functional parallels to animal and human immunity and thus the study of bacterial antiphage defense can yield insights into human innate immunity^7–9^. Furthermore, exploitation of bacterial antiphage systems has inspired essential tools for molecular biology and biotechnology, from restriction enzymes that revolutionized molecular cloning^10^, to CRISPR which transformed gene editing^11^. Another impact of the study of antiphage defense system lies in phage therapy, given that defense systems likely limit the utility of phages for treatment.

Due to the biotechnological implications of the antiphage defense systems that have been previously characterized, much recent research has attempted to discover novel systems through bioinformatic analyses followed by experimental validation, often in non-native hosts. Because antiphage defense systems often co-localize into genomic defense islands, bioinformatic discovery approaches that use such co-localization to identify defense systems have been developed^6,12–15^. An alternative strategy that has not been nearly as widely employed involves manipulating immune islands directly in native hosts to identify active defense systems and determine their roles in phage tropism and regulation under native conditions^16,17^.

The restriction modification (RM) systems are a major group of phage defense systems that can restrict phage and plasmid replication by recognizing and cleaving either unmodified or, conversely, modified DNA. RM systems are grouped into four types based on their mechanism by which the substrate is targeted^18,19^. Type II RM systems are the most common^20^ and typically include a modification methyltransferase that methylates ‘self’ DNA, while the restriction endonuclease cuts unmethylated, ‘foreign’ DNA either at the recognition site, or a few bases away^19,21,22^. In contrast, type IV RM systems recognize and restrict specific hypermodified ‘foreign’ DNA, whereas the ‘self’ DNA is not hypermodified and therefore is not restricted^23,24^. Type IV restriction systems, such as GmrSD and SauUSI, are also known as modification dependent restriction endonucleases (MDREs) and lack a methylase gene^1,25^.

MDREs are modular, containing a modification sensor domain and an effector nuclease domain. There is a wide range of sensor domains, of which the most common ones belong to the PseudoUridine synthase and Archaeosine transglycosylase (PUA) superfamily, including YTH, SRA, EVE and ASCH domains, all of which are thought to recognize specific types of modification^24,26–28^. Recently, a winged helix sensor domain was described^29^, and there is likely a larger variety of modification sensor domains that are yet undiscovered. The nuclease domains of MDREs typically belong to the PD-DExK, GIY-YIG, HNH and phospholipase D-like (PLD) superfamilies, each of which has specific cleavage requirements, such as presence of divalent cations^21,29,30^.

The incessant evolutionary arms race between phages and bacteria has produced a complex interaction network between multiple bacterial defense systems and phage-encoded inhibitors which impart immunity to these host systems^31,32^. For example, phage T4, which contains hypermodified cytosine in its genome, encodes multiple small-protein inhibitors of GmrSD and closely related endonucleases delivered during the DNA entry stage of infection^33,34^.

Here, we identify a single-gene defense system in a clinical *P. aeruginosa* strain CF040 which we call END^PaCF1^ (see below). Unlike previously characterized Type IV modification-dependent restriction endonucleases, END^PaCF1^ appears to be agnostic to the exact modification of the foreign DNA and can target DNA molecules containing a wide variety of hypermodifications occurring on both pyrimidines and purines. We attributed this ability to broad recognize a wide range of phages with hypermodified DNA to the C-terminal domain (CTD) of END^PaCF1^ which is homologous to the eukaryotic Ogg1 DNA glycosylase and prokaryotic Endonuclease III, enzymes that detect and excise non-canonical bases. Comparative genomic analysis shows that END^PaCF1^ belongs to a larger group of nucleases with inactivated **<u>En</u>**doIII **<u>d</u>**omains (END) that are predicted to recognize a broad range of hypermodified bases. Using END^PaCF1^ we show that Pbunaviruses, a well-studied family of phages commonly used for *P. aeruginosa* phage therapy^35^, have hypermodified genomes. Furthermore, some Pbunaviruses encode an inhibitor that is necessary and sufficient for END^PaCF1^ inhibition.

## Results

### Cascade Cas3 can be used to interrogate large defense islands to identify endogenously functional defense systems

Cascade-Cas3 has previously been developed to delete large parts of the *P. aeruginosa* chromosome^36^. We adapted the all-in-one vector carrying Cascade-Cas3 and the guide to delete large defense islands in *P. aeruginosa* clinical isolates. The clinical isolate CF040 was selected as it harbors two predicted defense islands which include well-characterized Type I RM systems (*hsdMSR*)^37^. Therefore, guides against the *hsdM/R* genes of each island were designed, introduced into the all-in-one vector and used to transform CF040. Transformants were then grown in the presence of gentamicin and the rhamnose inducer. After targeting, two surviving colonies with putative deletions in the same defense island were selected (τ<*di1*-1 and τ<*di1*-2), and whole genome sequencing revealed that τ<*di1*-1 (29kb) and τ<*di1*-2 (36kb) had different parts of defense island 2 deleted (Figure 1a; see Extended Data Table 1 for gene annotation). Plaque assays using these deletion strains showed that while the Pbunavirus phages F8 and PB-1 did not plaque well on CF040 and CF040τ<*di1*-1, there was a three-log increase in plaquing between clones CF040τ<*di1*-1 and CF040τ<*di1*-2 (Figure 1b). This finding suggests that clone CF040τ<*di1*-1 encodes a defense system(s) against the Pbunavirus phages that was deleted in clone CF040τ<*di1*-2. BLASTP searches of non-redundant (nr) protein sequences at the NCBI with each of the proteins encoded by the four genes deleted in CF040τ<*di1*-2 as queries showed that one of these proteins was annotated to contain a “PLDc-BfiI DEXD like” region (CDD: 197216) that is characteristic of type IIS restriction nucleases. Neither DefenseFinder nor PADLOC identified this gene as a defense system. Independent deletion of the gene encoding this putative type IIS-related enzyme showed that this single gene was responsible for inhibiting the *Pbunavirus* phages in CF040 (Figure 1b).

**Figure 1:**
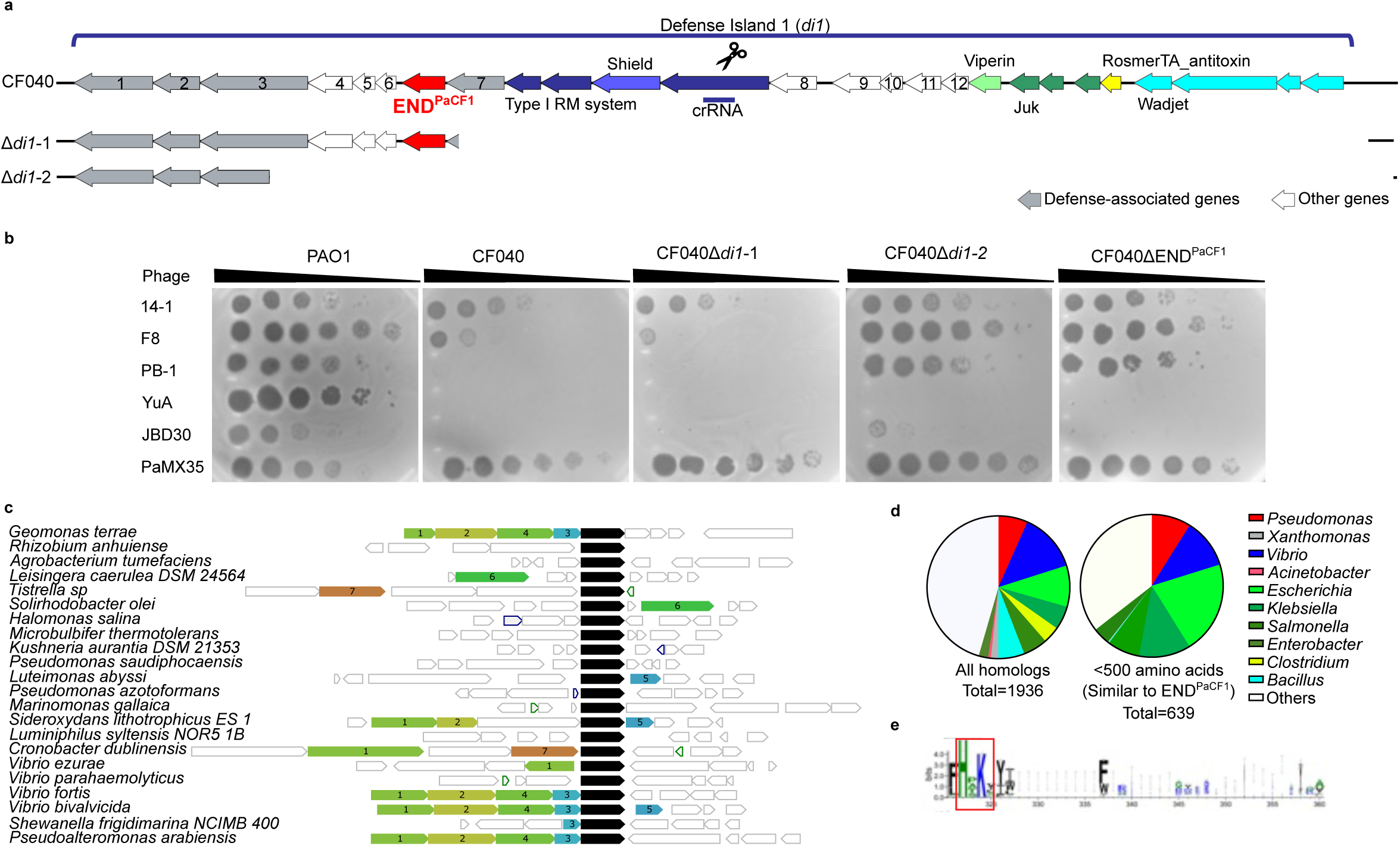
Identification of type IIS-like endonuclease END^PaCF1^ in clinical isolate CF040 using CASCADE Cas3 technology. **a**, Deletions generated in clinical isolate CF040 using CASCADE Cas3 against each type I restriction modification (RM) system in each of the two putative islands found in CF040. Grey arrows: Defense-related genes; white arrows: other genes (see Extended Data Table 1 for annotation of each numbered gene) The Type-IIS-like endonuclease was renamed as END^PaCF1^. The Cascade Cas3 guide designed against the *hsdR* gene in this defense island is indicated as crRNA. **b**, Plaque assays with different phages onto CF040 with different genes within the immune island deleted (as seen in a). Phages are spotted in 10-fold serial dilutions onto bacterial lawns, and clearings represent phage replication. **c**, Amino acid sequence of END^PaCF1^ was input into web-FLAGS for neighbourhood analysis in other bacteria. 1: DNA methylase family; 2: Restriction endonuclease subunit S; 3: hypothetical protein; 4: AAA ATPase; 5: Restriction endonuclease; 6: VapE family protein; 7: BrxL. **d**, Genus of bacteria in which homologs of END^PaCF1^ are identified are shown in pie-chart. Homologs were shortlisted after three iterations of PSI BLAST, with a cut off of >70% coverage and E values < 0.0005. Left: all homologs; right: homologs with a similar size to END^PaCF1^ (i.e. < 500 amino acid). **e**, Consensus sequence of amino acid near the endonuclease active site (HxK, boxed in red). Sequences (n=639) aligned as per previous were input into WebLogo3. The height of the amino acid lettering corresponds to the conservation of this amino acid at the indicated position.

Thus, the all-in-one Cascade-Cas3 vector can be employed to efficiently interrogate defense islands in clinical isolates, and uncovered a novel, single-gene defense system which we name END^PaCF1^.

### END^PaCF1^ is an endonuclease represented in diverse bacteria

Restriction modification (RM) systems often consist of a methylase gene adjacent to the endonuclease, the function of which is to help distinguish self-DNA from non-self^19^. Therefore, to determine if END^PaCF1^ is associated with a methylase, we analyzed gene neighborhoods using WebFlaGs^38^ and found that the gene encoding END^PaCF1^ is in defense islands in various bacteria, but no other gene appeared to be co-associated with it across genomes (Figure 1c). In particular, no adjacent gene encoding a methyltransferase was found, suggesting that this is not a canonical RM system.

A PSI-BLAST search of the non-redundant protein database using END^PaCF1^ as the query identified homologs of this protein in numerous bacteria, both clinical pathogens and environmental microbes (Figure 1d). All END^PaCF1^ homologs contained the active site HxK motif essential for catalysis of phosphodiester backbone cleavage, similar to type IIS enzymes BfiI and NgoAVII (Figure 1e)^39,40^. All of these endonucleases belong to the phospholipase D (PLD) superfamily of hydrolases that cleave diverse substrates including the phosphodiester bonds of dsDNA^30,41^. However, BfiI and NgoAVII are co-encoded with methyltransferases in canonical Type IIS R-M systems^22^.

### END^PaCF1^ acts early in infection, is non-abortive, and targets diverse phage families

To determine which families of phages are restricted by END^PaCF1^, END^PaCF1^ was first introduced into the phage sensitive lab strain PAO1, which lacks an END^PaCF1^ homolog, on an episomal vector (pHERD30T, shortened to p30T) for overexpression or in the chromosome at the *att*Tn7 site^42^, under an inducible promoter. We challenged PAO1 p30T^END-PaCF1^ with a panel of diverse *P. aeruginosa* phages. Plaque assays with this overexpression strain showed that END^PaCF1^ inhibited growth of Yuavirus (YuA-like), Abidjanvirus (Ab18-like), Pbunavirus (F8 family of phages) and Wroclawvirus (Pa5Oct related jumbo phages) (Extended Data Figure 1a).

We next confirmed that PAO1 *att*Tn7::END^PaCF1^ also targeted all four phage families when induced (Extended Data Figure 1b). However, in the absence of induction, PAO1 *att*Tn7::END^PaCF1^ targeted F8 but not 14-1 (Figure 2a and Extended Data Figure 1b), which was similarly observed in CF040 (Extended Data Figure 1c). This finding suggests that expression of END^PaCF1^ in uninduced PAO1 *att*Tn7::END^PaCF1^ approximates the expression level in the native CF040 strain. END^PaCF1-dead^, in which the essential catalytic residues of the HPK active sites were replaced with alanine (APA), failed to confer protection against phages, confirming that phage targeting is due to the endonuclease activity of END^PaCF1^ (Extended Data Figure 1b). Of note, no escapers for phages F8, PB-1, YuA, PaMX11 and Ab18 were isolated on PAO1 *att*Tn7::END^PaCF1^, suggesting that phages cannot easily escape the system (≥10^5^ phages in the first spot) (Extended Data Figure 1b).

**Figure 2:**
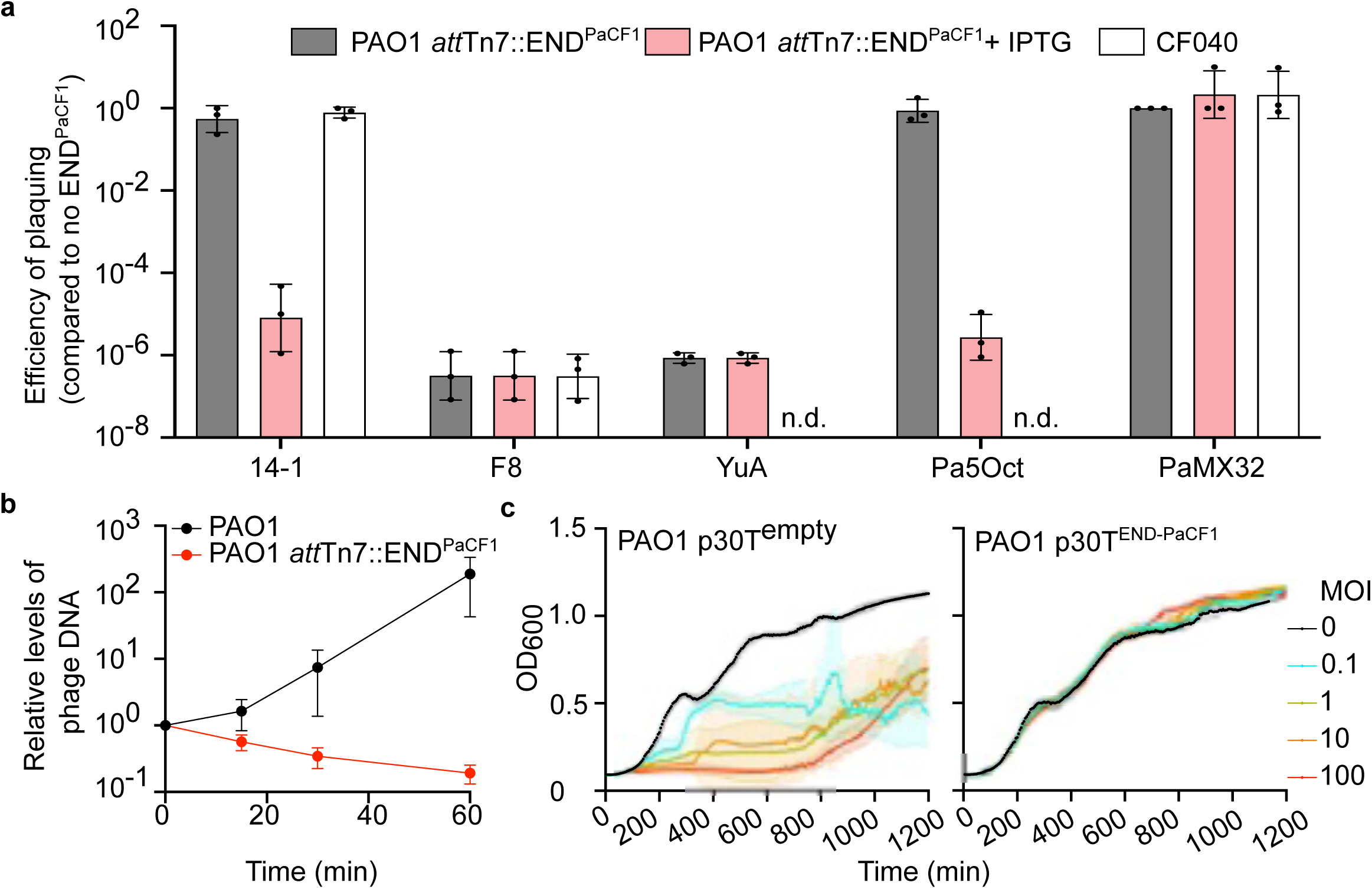
END^PaCF1^ functions within the half hour of infection and is not abortive. **a**, F8-like phages 14-1 and F8, YuA and Pa5Oct were plaqued onto PAO1 and onto PAO1 carrying END^PaCF1^ on the chromosome at the Tn7 integration site and CF040 with/without END^PaCF1^. Efficiency of plaquing is calculated as percentage with respect to phage titer on PAO1/CF040ΔEND^PaCF1^. PaMX32 is included as a plaque control, n.d. refers to not detected. 1mM IPTG was included to induce expression of integrated END^PaCF1^. **b**, qPCR time course assay F8 infection in PAO1 and PAO1 *att*Tn7::END^PaCF1^ without induction. Samples (500 **μ**l) were taken at t= 0, 15, 30, and 60 minutes post infection. Values were first normalized to PAO1 gDNA using primers binding to *rpoD*, before normalizing to t = 0. **c**, Growth curves of PAO1 with/without END^PaCF1^ infected with F8 at various MOIs.

To determine whether END^PaCF1^ inhibits phage reproduction early or late in infection, PAO1 and PAO1 *att*Tn7::END^PaCF1^ were infected with F8 and the bacteria were collected every 15 min. qPCR results demonstrated that in the absence of END^PaCF1^, the F8 genome replicated approx. 100-fold over 60 minutes; however, no replication of the F8 genome was observed in PAO1 *att*Tn7::END^PaCF1^ (Figure 2b). Thus, END^PaCF1^ appears to act early in infection, antagonizing DNA replication.

Given that a broad variety of bacterial defense systems induce dormancy or programmed cell death (a phenomena often denoted as abortive infection)^43–47^, we sought to determine whether END^PaCF1^ was abortive. To this end, PAO1 was transformed with p30T with or without END^PaCF1^ to intentionally overexpress the system. Both strains were then infected with F8 at high multiplicity of infection (MOIs) and tracked over 20 hours. We observed no decrease in the bacterial growth rate at high MOIs and overexpression of END^PaCF1^ in the absence of phage did not decrease growth rate of PAO1 (Figure 2c). These results indicate that END^PaCF1^ is not an abortive infection system.

To test whether YuA (Yuavirus), Ab18 (Abidjanvirus), 14-1 and F8 (Pbunavirus) and Pa5Oct (Wroclawvirus) were also targeted by END^PaCF1^ homologs in other strains, BLASTP was used to identify strains carrying END^PaCF1^ homologs. The *P. aeruginosa* lab strain PAK was found to encode a homolog of END^PaCF1^, END^PAK^, with 98% amino acid identity. When END^PAK^ was deleted from their respective native strains, F8 and PB-1 phages formed plaques. There were no visible changes upon infection with Yuavirus, Abidjanvirus and Pa5Oct, suggesting that these phages were blocked through another mechanism (Extended Data Figure 1c).

Additional homologs with different levels of amino acid identity as determined by BLASTP were introduced into PAO1 (see Extended Data Table 2 for full list of homologs tested). Plaque assays showed that END^PaCF1^ homologs END^Pa3^ (91% amino acid identity to END^PaCF1^ from *P. aeruginosa*) and END^Pa4^ (40% amino acid identity to END^PaCF1^ from *P. aeruginosa*) caused a 4-5 log reduction for phages 14-1, F8, YuA, Ab18 and Pa5Oct. In contrast, END^Se^ from *Salmonella enterica* (50% amino acid identity) resulted in a 3-log reduction for phages Ab18 and Pa5Oct (Extended Data Figure 1d). These results confirm that the family of END^PaCF1^ nucleases can target phages of these distinct families.

### END homologs target diverse hypermodified phages

Of the targeted phages, Yuavirus M6 carries 5-(2-aminoethyl) uridine (5-NedU) instead of thymine in its genome^48^. The closely related Abidjanvirus Ab18 is predicted to also contain a hypermodified thymine^40,41^. However, the genomes of the other targeted phages targeted by END^PaCF1^ were not known to be hypermodified. To detect potential modifications, F8 and Pa5Oct genomes were subjected to *in vitro* digestion by various commercially available restriction enzymes with well-defined site specificities (Extended Data Table 3). Pa5Oct, a jumbo phage unrelated to phiKZ-nucleus forming phages^49^, was resistant to EcoRI, BmrI, ApaLI, XbaI and BsaI, suggesting that it contains a modified G, in a GA and/or GG motif (Extended Data Table 2, Figure 3a). Analysis of the Pa5Oct genome with Domainator^50^ followed by confirmation with Foldseek^51^ also revealed genes associated with biosynthesis of 7-deaza-2’-deoxy-G such as gp162 (QueE; E-value 3.3 E^−12^) and gp230 (6-pyruvoyl tetrahydropterin synthase; E-value 5.5 E^−14^), further suggesting that Pa5Oct has a modified G.

**Figure 3:**
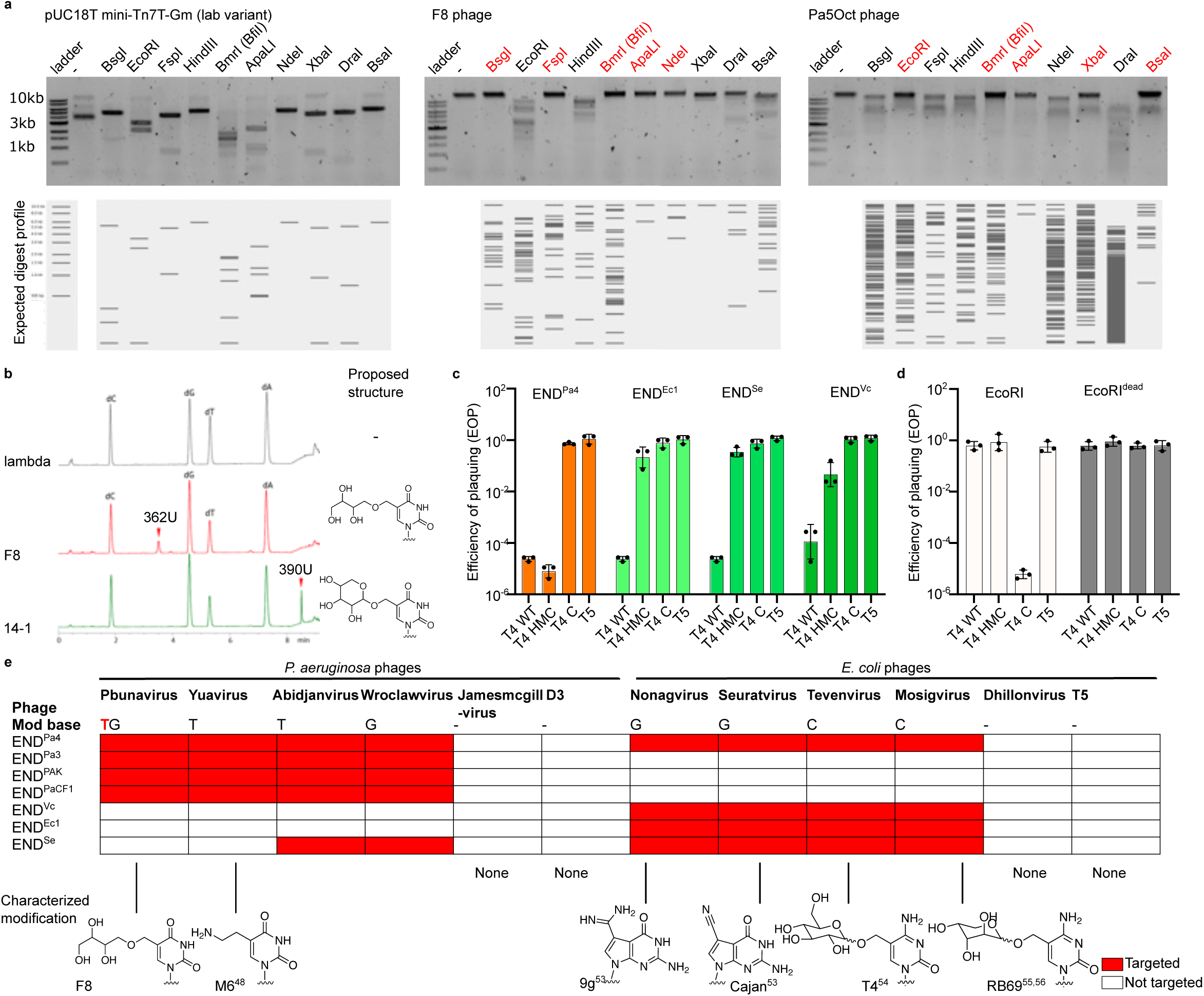
Pbunavirus F8 and 14-1 has a modified thymidine in its genome; END^PaCF1^ therefore targets only phages with modified genomes, but is agnostic to the exact modification. **a**, Restriction digest of pUC18T mini-Tn7T-Gm plasmid (lab variant), F8 genome and Pa5Oct genome with restriction enzymes. Restriction enzymes in red indicate that they did not digest F8/Pa5Oct phage shown. **b**, Mass spectrometry of F8 and 14-1 genomic DNA. Lambda phage, which does not have a modified genome, is included as a control. **c**, T4 phages with wildtype, hydroxylmethylcytosine (HMC) or unmodified cytosine (C) were plaqued against p30T carrying homologs of END^PaCF1^, as indicated on the graph. T5 was included as a control. Efficiency of plaquing shown is relative to empty vector. **d**, T4 phages with different modifications were plaqued against DH10B with functional or dead EcoRI. Efficiency of plaquing shown is relative to empty vector; T5 was included as a control. **e**, Table of phages that are targeted by different homologs of END^PaCF1^. An example of phage DNA hypermodifications is shown, including the phage from which the characterization was done. Only the F8 hypermodification is described here; the others were obtained from previously published papers.

We presumed that F8 phage contained no modifications because we have previously demonstrated digestion of its DNA *in vivo* with EcoRI^52^. However, we found that, of the ten restriction enzymes tested, only EcoRI, HindIII, XbaI, DraI, BsaI digested F8 DNA (Figure 3a). Virion DNA from phage F8 is not cleaved by BsgI, FspI, BmrI, ApaLI and NdeI. The enzymes which could not digest F8 all contain a TG dinucleotide within their recognition sequences, which is not present in the recognition sites of EcoRI, HindIII, XbaI, DraI, BsaI (Extended Data Table 3).

To identify the modified base, virion DNA purified phages F8 and 14-1 were enzymatically digested to free nucleosides and subjected to high-performance liquid chromatography/mass spectrometry (LC-MS). The chromatograms of F8 and 14-1 nucleoside digests revealed a novel nucleoside peak for each, with an apparent mass of 362 U and 390 U, respectively. Quantitation of peaks from canonical nucleotides shows a T:A ratio of less than one, indicating that these novel nucleosides to be thymidine derivatives replacing approximately one quarter of the canonical thymidines. The observed masses and the follow-up fragmentation analysis are consistent with an erythrose/threose sugar alcohol derivative in phage F8, and a circularized aldopentose derivative in phage 14-1 (Figure 3b, Extended Data Figure 2). Taken together, these restriction digests and analytical methods suggest that the Pbunavirus DNA previously thought to be unmodified, contains hypermodified thymine, in a TG motif. This confirms that the END nucleases appear to only target phages with hypermodified genomes, and that they can be used to reveal that a phage has a hypermodified genome when it is not previously known.

To determine if END nuclease homologs also recognized and/or targeted hypermodified DNA, previously identified homologs (END^Pa3^, END^Pa4^, END^Se^) and additional homologs identified in other Gram-negative bacteria were expressed in *Escherichia coli* BW25113 for plaque assays. These assays confirmed that the *E. coli* homologs of END^PaCF1^, specifically END^Vc^, END^Ec1^ and END^Se1^, targeted all hypermodified phages tested in *E. coli*, but not T5-like phages (Extended Data Figure 3 and Extended Data Table 2 for list of homologs). These hypermodified coliphages included *Nonagvirus*^53^, *Seuratvirus*^53^, T-even phages^54^ and Mosigvirus^55,56^, with a total of at least four different hypermodifications over two bases (G and C). T5 and related phages were used as a control as they lack hypermodifications in their genomes.

To further confirm that the phages were targeted due to the presence of hypermodification in the genome, we tested targeting of wildtype T4 containing glucose-hydroxymethyl-cytosine, T4 containing hydroxymethyl-cytosine (T4-HMC) and T4 with an unmodified genome (T4-C) in an *E. coli* DH10B strain to maintain the phage DNA modifications. The same phages were then used to infect DH10B carrying different END homologs. In this experiment, END^Pa4^ restricted both the wildtype T4 and T4-HMC but not T4-C. In contrast, homologs from *E. coli*, END^Ec1^ and the closely related END^Se^, caused a five-log decrease of plaquing efficiency for wildtype T4 but had modest protection against T4-HMC (Figure 3c), which is similar to the protection profile of type IV restriction endonuclease GmrSD^25^. END^Vc^ caused a four-log decrease in plaquing efficiency for T4 and a one-log decrease in plaquing efficiency of T4-HMC phage (Figure 3c). No END homologs restricted T4-C, which was in turn restricted by EcoRI as EcoRI targets non-modified DNA (Figure 3d)^57^. This confirms that END nuclease homologs recognize hypermodified DNA.

Overall, END^Pa4^ was found to target eight distinct hypermodified phage families with at least ten different known hypermodifications. Similarly, END^Se^ targeted all known modified phages in *E. coli*, as well as two groups of hypermodified phages in *P. aeruginosa*, Abidjanvirus and Wrowclawvirus. Other *P. aeruginosa* homologs, (END^PAK^, END^Pa3^, END^PaCF1^) targeted all known hypermodified phages in *P. aeruginosa*, whereas END^Ec1^ and END^Vc^ targeted hypermodified phages in *E. coli*. Thus, all END homologs each targeted multiple hypermodifications (Figure 3e). None of the END homologs tested restricted growth of phages lacking hypermodified bases in their genomes in either *P. aeruginosa* or *E. coli* (Figure 3e). These results confirm that END homologs target only phages with hypermodified genomes, with some homologs targeting regardless of the specific nature of the hypermodification.

### C-terminal domain of END^PaCF1^ determines substrate specificity and defines a larger group of END nucleases

We showed that END nucleases define a distinct family of promiscuous MDREs that appeared to be agnostic to which hypermodification they target (Figure 3e). We therefore sought to determine which domain was responsible for detecting the hypermodified genomes. HHpred analysis^58^ revealed significant similarity of END^PaCF1^ CTD with endonuclease III family members, in particular, that from *Deinococcus radiodurans* (DrEndoIII) and eukaryotic Ogg1 DNA glycosylase (including human hOGG1) (Figure 4a, Extended Data Figure 4). Comparison of the AlphaFold3^59^ structural model of END^PaCF1^ with the structures in the Protein DataBank confirmed similarity between the NTD and PLD superfamily enzymes and between CTD and ENDOIII / hOGG1 (Figure 4a), but not to the predicted structure of the Brig1 DNA glycosylase defense system recently characterized^60^ (Figure 4b). DrEndoIII detects and targets oxidized T and C bases for base excision repair in prokaryotes, whereas hOgg1 excises 8-oxo-7,8-dihydro-2’-deoxyguanosine in mammalian cells^61–63^. END^PaCF1^ lacks the catalytic residues of the endonuclease III family and therefore is likely to be enzymatically inactive (iEndoIII) (Extended Data Figure 4b). We therefore hypothesized that the CTD hOgg1/iEndoIII domain of END^PaCF1^ is responsible for detecting hypermodified genomes by recognizing the respective non-canonical bases. We first confirmed the conservation of the predicted structure and orientation of the CTD in the more distant homologs, END^Pa4^ and END^Ec1^ (approx. 40-50% amino acid identity to END^PaCF1^ and to each other) (Figure 4c). Next, to determine if CTD determines specificity, END^Pa4^ endonuclease was fused END^Ec1^ CTD, resulting in fusion protein END^Pa4-Ec1-fusion^ (Figure 4d). All three nucleases END^Pa4^, END^Ec1^ and END^Pa4-Ec1-fusion^ fully targeted phages Bas21 and Bas47. However, similar to END^Ec1^ but not END^Pa4^, END^Pa4-Ec1-fusion^ preferentially targets T4 glu-HMC over T4-HMC (Figure 4e, Extended Data Figure 5). This result suggests that the CTD iEndoIII domain could be responsible for the specificity/substrate selection by END^PaCF1^.

**Figure 4:**
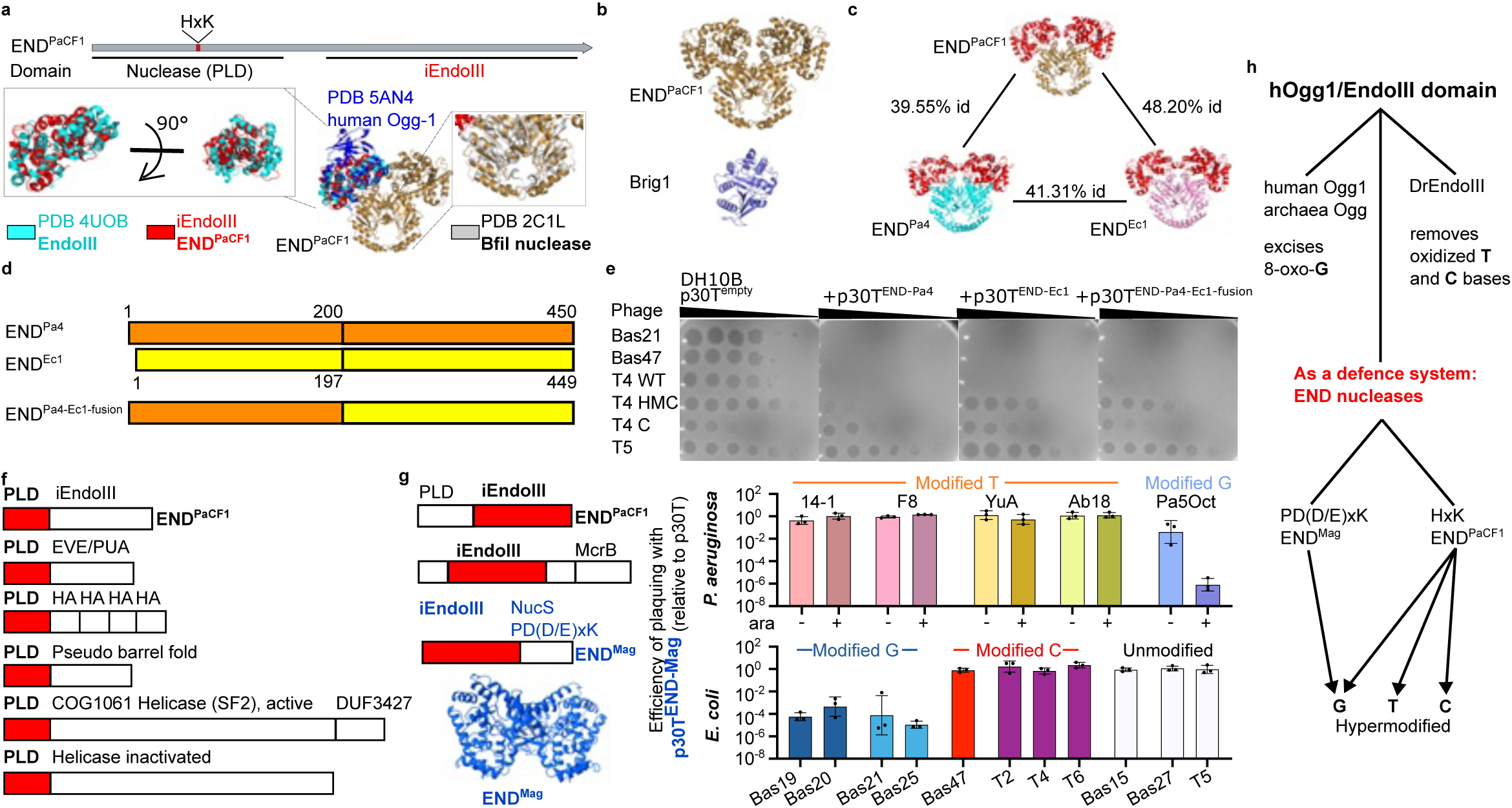
Domain organization and function of END nucleases. **a**, Domain organization and AlphaFold model for END^PaCF1^. Approximately domain boundaries are based on HHpred results (Extended Data Figure 4). The N-terminal nuclease domain belongs to phospholipase D-like (PLD) family. Bottom: Overlay (cealign) of Alphafold3 prediction of END^PaCF1^ domains with known structures PDB 4UOB (endonuclease III, EndoIII), 5AN4 (human Ogg-1) and 2C1L (BfiI) **b**, AlphaFold model for END^PaCF1^ compared to the prediction for Brig1. **c**, AlphaFold model for END^PaCF1^ and distant homologs. The percent amino acid identity between each homolog is as shown. The C-terminal domain (CTD) is shown in red. **d**, Schematic of swap between CTD from END^Pa4^ homolog and END^Ec1^ homolog. The amino acids and the numbered position indicate position of swap. **e**, Plaque assays of Bas21, Bas 47 and T4 phages carrying different modifications on its genome with END^Pa4^, END^Ec1^ and END^Pa4-Ec1-fusion^ shown in d. HMC = hydroxymethylcytosine, C = unmodified cytosine. Plaque assays are representative of three repeats. **f**, Domain organization of other HxK/PLD nuclease fusions. The nuclease domain is colored red. Abbreviations: PUA - pseudouridine synthase and archaeosine transglycosylase domain, same family as EVE domain, HA -helicase associated domain, COG - cluster of orthologous genes, DUF - domain of unknown function. **g**, Domain organization of other of iENDO fusions. The iEndoIII domain is colored red. McrB is a GTPase associated with type IV restriction-modification systems and NucS belong to PD(D/E)xK nuclease superfamily. END^Mag^ (Alphafold structure prediction shown in blue), was tested in both *P. aeruginosa* (top) and *E. coli* (bottom). For *P. aeruginosa*, the system is further induced with arabinose (ara) **h**, Schematic of targets of known proteins carrying the hOgg1/EndoIII domain.

To further determine if specificity is determined by the CTD, we sought to identify other endonucleases containing a homologous PLD NTDs, but with different CTDs. Indeed, a PSI-Blast search identified PLD nucleases fused to several unrelated CTDs including EVE (of the PUA superfamily), helicases, helicase-associated domains previously shown to bind DNA^64^, and pseudobarrel domains (Figure 4f). The NTD PLD nuclease fused to CTD pseudo-barrel or helicase domains have been identified as functional restriction enzymes BfiI and SauUSI, respectively^1,40^. Consistent with our hypothesis that substrate specificity is due to the CTD, SauUSI targets 5-methylcytosine DNA^1^ whereas BfiI is encoded next to a methyltransferase and targets unmodified DNA^22,40^. SauUSI, in particular, is capable of restricting T4 phage infection^1^. Thus, END^PaCF1^ belongs to a larger group of defense endonucleases featuring an N-terminal PLD nuclease and a variable C-terminal modification-recognition domain.

In a reciprocal search, we identified the iEndoIII sensor fused to two distinct catalytic domains, a “NucS”-like PD-DExK superfamily nuclease, and the McrB-like GTPase. The PD-DExK-iEndoIII homolog from a metagenome (Accession MDV2503288 “END^Mag^”), showed high similarity to archaeal Ogg1 enzymes that recognizes 8-oxo-guanine, similar to hOgg1^65^, as demonstrated by HHpred search (Extended Data Figure 6). We cloned END^Mag^ into PAO1 and *E. coli* BW25113 and found that, in plaque assays END^Mag^, like END^PaCF1^, targeted phages with modified G (Pa5Oct, Bas19, Bas20, Bas21, Bas25) but not phages with hypermodification on T or C or phages with unmodified genomes (Figure 4g). Thus, whereas Ogg1 and EndoIII target oxidized G and T/C bases respectively, END^PaCF1^ and END^Mag^ co-opted the inactivated hOgg1/iEndoIII domain as a sensor of modified bases to target hypermodified phage genomes (Figure 4h). Given that i**<u>En</u>**doIII **<u>d</u>**omain of these endonucleases appears to be responsible for their ability to recognize diverse hypermodified phage genomes, we propose to denote this group of defense enzymes collectively as **<u>END</u>**nucleases.

### A single anti-END phage protein inhibits diverse END nuclease homologs

We previously observed that F8 and PB-1 were targeted by END^PaCF1^ in CF040, 14-1 was not (Figure 1c), but 14-1 is highly similar to F8 and PB-1 according to a gene cluster comparison (Figure 5a). To better understand what 14-1 gene facilitates escape from END^PaCF1^, we made hybrid phages of 14-1 and F8 by co-infecting PAO1 with both phages followed by selection against parental 14-1 and F8 using Type I-C Cascade-Cas3 and END^PaCF1^ respectively. Analysis of the assembled hybrid phage genomes suggested three candidate regions that could harbor potential inhibitors. Sequences within each region were then individually queried for 14-1 and F8 SNPs across all hybrid phage genomes. It was found that all hybrid phages carried gp88 from 14-1, suggesting that this protein is an inhibitor of END^PaCF1^ (Figure 5b).

**Figure 5:**
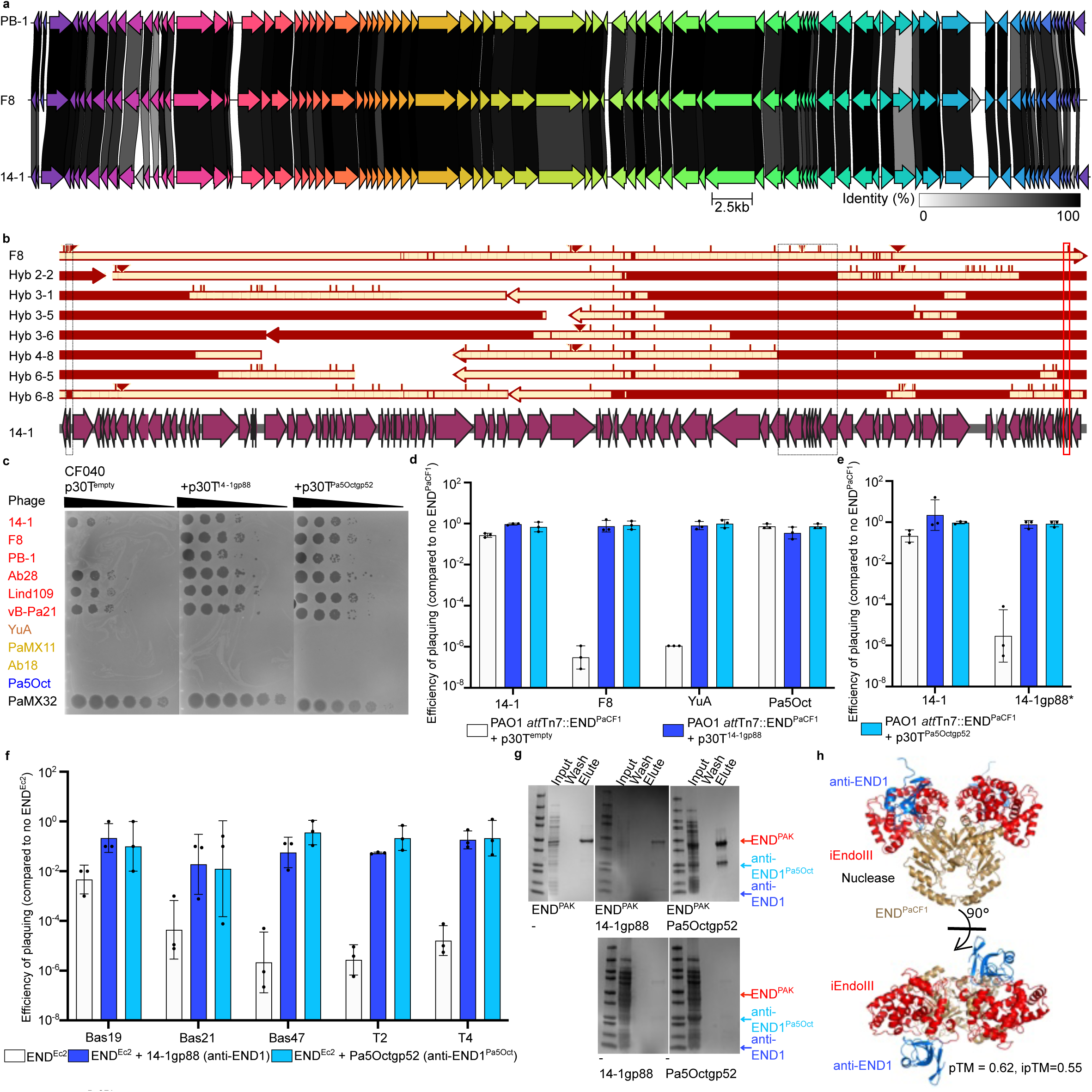
END^PaCF1^ inhibitors are encoded by some Pbunavirus phages including 14-1, as well as unrelated phage Pa5Oct and cross protect targeted phages from other families. **a**, Alignment of F8, 14-1 and PB-1. 14-1 is not inhibited by END^PaCF1^ in CF040, while F8 and PB-1 are. Alignment was done with clinker. **b**, Alignment of hybrid phages of 14-1 and F8. Hybrid phages of 14-1 (not targeted by endogenous END^PaCF1^ in CF040) and F8 (targeted by endogenous END^PaCF1^ in CF040) were constructed by co-infection, then sequenced with Illumina sequencing prior to assembly using SPADES. Alignment on Snapgene (shown here) and parsing with grep revealed all hybrid phages carried 14-1 gp88 (shown with a red box). **c**, Panel of *P. aeruginosa* phages were tested against END^PaCF1^ in the native strain CF040 with inhibitors cloned into p30T plasmid. Plaque assay shown is representative of three repeats. **d,** Plaque assay with PAO1 carrying END^PaCF1^ integrated into *att*Tn7 site, with 14-1 gp88 or Pa5Oct 0052 expressed in *trans*. Empty p30T plasmid was included as a control. **e**, Plaque assay with wildtype 14-1 or 14-1 with gp88 (14-1gp88*) with a frameshift mutation, onto PAO1 *att*Tn7::END^PaCF1^. Presence of either inhibitor rescued plaquing of 14-1gp88*. **f,** Plaque assay with different hypermodified phages in *E. coli*, in the presence of END^Ec2^ with/without 14-1 gp88 (anti-END1) or Pa5Oct gp52 (anti-END1^Pa5Oct^) inhibitors on pDUET. **g**, Pulldown of inhibitors in the presence of defense system. Only END^PAK^ is His-tagged. **h**, Alphafold3 prediction of END^PaCF1^-anti-END1 interaction. The predicted structure shown is consistent across all five predicted outputs.

To test the prediction that 14-1 gp88 inhibits END^PaCF1^, gp88 and its homolog identified using BLASTp, gp52 from jumbo phage Pa5Oct, were introduced into the native strain CF040. Expression of 14-1 gp88 or Pa5Oct gp52 increased F8 and PB-1 plaquing efficiency by five logs compared to wildtype. No difference in plaquing efficiency was observed for 14-1 (Figure 5c). To confirm that this effect was due to the inhibition of END^PaCF1^ and not other immune systems present in CF040, both inhibitors were expressed in the presence of END^PaCF1^ in PAO1 and facilitated plaquing of all families of phages. These results suggest that the inhibitor is not phage-family nor hypermodification specific (Figure 5d). Furthermore, Cascade-Cas3 targeting of gp88 enabled the isolation of a 14-1 phage mutant with an early stop codon in gp88 due to a frameshift mutation (14-1^gp88*^). Compared to 14-1, 14-1^gp88*^ was susceptible to END^PaCF1^ in the native host but was rescued by either of the END^PaCF1^ inhibitors expressed *in trans* (Figure 5e). Additionally, these phage proteins also inhibited the *E. coli* homolog END^Ec2^ (Figure 5f). Therefore, both 14-1 gp88 (anti-END1) and Pa5Oct gp52 (anti-END1^Pa5Oct^) blocks multiple END homologs regardless of the exact DNA hypermodification targeted by END nucleases.

To test whether the identified inhibitors specifically bound END^PaCF1^, anti-END1 and FLAG-tagged anti-END1^Pa5Oct^ were cloned together with His-tagged END^PAK^ (90% identity to END^PaCF1^) into the pETDuet-1 vector in *E. coli* BL21 and were shown to co-purify on a Ni-NTA column (Figure 5g). Additionally, we showed that both inhibitors co-purified with END^Pa4^, a distant homolog of END^PaCF1^ in *P. aeruginosa* (40% amino acid identity), and inhibited END^Pa4^ *in vivo* in *E. coli* (Extended Data Figure 7). An AlphaFold3^59^ prediction of the defense system-inhibitor complex also suggests with modest confidence that anti-END1 binds END^PaCF1^ (pTM = 0.62, ipTM=0.55) in the iEndoIII region (Figure 5h).

Overall, we have identified a group of nucleases, the END nucleases, that is causal for phage infection failure in a clinical isolate and target multiple DNA hypermodifications. To counter the END nucleases, some phages with hypermodified genomes encode anti-END inhibitors that can block multiple END homologs and bind directly to these nucleases.

## Discussion

The recent discovery of an unexpected, unprecedented diversity of antivirus defense systems in prokaryotes sparked much interest in prokaryotic immunity and stimulated the development of computational and experimental approaches and tools to identify immune systems and defense islands^5,6,12–14^. However, few tools exist to manipulate these islands in clinical strains, and over multiple genes. In this work, we adapted the recently developed Cascade-Cas3 technology for two purposes: first, to generate large deletions in defense islands, resulting in the identification of a distinct, single-gene defense system, and second, to probe phage genomes for inhibitors of defense systems. Given that the Cascade-Cas3 vector has been shown to be functional in other species, this approach can be widely exploited to interrogate defense islands and phages of other bacteria species in their native conditions.

The single-gene defense system identified in this work is an endonuclease homologous to type IIS endonucleases which belong to the PLD hydrolase superfamily. However, unlike previously studied type IIS endonucleases^1,25,29^, END^PaCF1^ recognizes hypermodified DNA, without discriminating between different types of hypermodification including modifications with and without sugar moieties, and modifications on different nucleotide bases. We proposed that the promiscuous recognition of hypermodified bases is mediated by the C-terminal domain of END^PaCF1^, an inactivated endonuclease III (iEndoIII). We found that iEndoIII domain can also combine with other nucleases that are predicted to target hypermodified DNA as well. Similarly to other domains involved in multiple defense systems, such as TIR and Sir2^66^, the EndoIII domain is present across all three domains of life, eukaryotes^61^, bacteria^62^ and archaea^65^. The EndoIII domain is part of a larger helix-hairpin-helix (HhH) DNA glycosylase superfamily implicated in sensing damaged DNA for repair^65,67,68^. Mechanistic study of the HhH fold suggests that it can be easily mutated and evolved to recognize DNA adducts across all domains of life^69–71^. We show here that the inactivated EndoIII/Ogg derivative instead now serves as a sensor for multiple types of non-canonical bases in phage defense systems. We thus coined the term END nucleases for the defense enzyme with this type of modular organization.

The END nucleases are present in a broad variety of bacteria, from human pathogens such as *P. aeruginosa*, *E. coli*, *Klebsiella pneumoniae* to diverse environmental bacteria such as *Rhizobium*, *Xanthomonas* sp. and probably many more as suggested by the identification of these genes in metagenomes. These findings imply that phages with hypermodified genomes are more common than currently known, and the vast array of DNA hypermodification mechanisms in phages and the defense systems targeting hypermodified DNA with varying degree of specificity remain to be characterized.

Additionally, using the END nucleases, we have identified the presence of hypermodified genomes on two phage families which were not known to be hypermodified previously. Currently, the identification of hypermodification in phage genomes is laborious, often relying on the presence of gene homologs previously implicated in modifying phage genomes, followed by confirmation with mass spectrometry analysis. However, as shown in this paper, this could miss out phages which do not carry such genes. One such example is the Pbunavirus family, which, despite being well studied due to its use in phage therapy, was not previously known to carry a hypermodified genome. Therefore, the END nucleases will provide a simple yet important tool to identify phages carrying a potentially hypermodified genome.

Our findings on END nucleases such as END^PaCF1^ support the modular model of modification dependent endonucleases^18^, where the NTD is responsible for the catalytic activity, whereas the CTD is the sensor. In accord with this model, we found that PLD superfamily catalytic domains homologous to that of END^PaCF1^’s also can combine with CTD sensors unrelated to iENDOIII, such as EVE, winged helix-turn-helix, pseudo barrel fold, helicases and helicases associated domains^1,22^, which likely confer distinct substrate specificities. The modularity observed among END and other defense endonucleases is consistent with the previous observations on MDREs^24^, antivirus STAND ATPases^18,44,72^, and type III CRISPR systems^73^ in all of which different sensors are combined with diverse effectors. This combinatorial modularity, driven by the Red Queen evolutionary dynamics, is emerging as a general principle of the evolution of immune systems. Another prominent aspect of the ubiquitous host-pathogen arms race is the evolution of defense system inhibitors by viruses, best explored in the case of anti-CRISPR proteins^74^. Here, we identified a family of anti-END inhibitors encoded by unrelated phages with distinct hypermodifications on DNA that protects against the END nuclease which *P. aeruginosa* phages are not able to escape from. A lack of escape is consistent with some phages being unable to simply lose their modified state, although this is possible for T4 and did induce END nuclease escape. Undoubtedly, many more combinations of prokaryotic immune systems with virus-encoded inhibitors or escape mechanisms remain to be discovered, and the study of their interactions will enrich our understanding of the strikingly complex landscape of microbial immunity.

## Methods

### Plasmid construction

To generate inserts in the pHERD30T (shortened to p30T) plasmid, the p30T backbone was amplified with primers WX_16/WX_17 (see Supplementary Table 1 for list of primers), before being subjected to DpnI (NEB) digest for 37 °C, 1 h. Inserts were amplified using primers with overhangs complementary to linearized p30T backbone, or ordered as gene fragments gblocks from Twist. For END nucleases from other bacteria, ATG was appended as the start codon if required. The inserts were then joined with the p30T backbone using Hi-Fi DNA Gibson Assembly (NEB) by incubating at 50 °C, 1 – 4 h. *E. coli* DH5α was then transformed with the resulting plasmids and verified by sequencing with Plasmidsaurus. *P. aeruginosa* cells were then electroporated with the p30T constructs and selected on 50 μg/ml gentamicin. p30T plasmids were also transformed into BW25113 and DH10B and selected on 15 μg/ml gentamicin. p30T sfCherry-EcoRI and sfCherry-EcoRI^dead^ have been previously constructed and tested for *in vivo* restriction function^57^ before transforming into DH10B.

To generate all-in-one Cascade Cas3 plasmids carrying different guides, plasmids carry the Cascade-Cas3 system and targeting sequences in the 14-1 phage or in the Type I RM system in CF040 were constructed by first digesting the empty vector with BsaI at 37 °C for 4 h. Guides with complementary overhangs (in the form of oligos ordered from IDT) were subjected to T4 Polynucleotide Kinase (NEB) treatment for 3 h, 37 °C and inactivated at 65 °C, 10 min. Guides were then annealed with their complementary strands by heating to 95 °C in a thermocycler for 10 minutes, then gradually ramping down to room temperature Guides were then diluted and ligated into BsaI-digested Cascade Cas3 with T4 ligase (Thermofisher Scientific) at rtp for 1h. DH5α were then transformed with the plasmid before recovering for 2h and selected on 15 μg/ml gentamicin. Cascade Cas3 plasmids with the correct guides (verified using Sanger sequencing) were then electroporated into PAO1/CF040.

To clone END^PaCF1^ and homologs into pETDuet, pETDuet with/without the inhibitor was digested with NdeI and XhoI for 4 h. Gene fragments carrying the defense system were amplified with the appropriate primers. Fragments were the ligated into the backbone using Hi-Fi Assembly according to manufacturer’s instructions, DH5α were then transformed with the ligated plasmid and then selected in the presence of 100 μg/ml carbenicillin. Colony PCR was then carried out with primers WX_216/WX_217; those with correct inserts were extracted and sent for sequencing with Plasmidsaurus or Quintara BioSciences. Desired plasmids were then transformed into BL21 or BW25113 using the heat shock chemical transformation protocol.

For cloning inhibitors into pETDuet, pETDuet with/without homologs of END^PaCF1^ was digested with NcoI and SalI. Gene fragments carrying the inhibitors were amplified with the appropriate fragments, ligated and selected as per previous. Colony PCR was carried out with primers WX_257/WX_258.

To insert END^PaCF1^ into the PAO1 chromosome, the mini-Tn7 insertion method was used^42^. Briefly, the defense system was first cloned into the lab’s mini-Tn7 plasmid. The plasmid backbone was first amplified with primers WX_129/WX_130; END^PaCF1^ and the corresponding 60bp upstream were amplified using primers WX_167/WX_168, before cloning into the plasmid. *P. aeruginosa* PAO1 cells were electroporated with the resulting plasmid and helper plasmid pTNS3 and selected for using gentamicin. The gentamicin marker was then excised with FLP recombinase as previously described^75^. Colonies were patched onto LB plates with/without gentamicin to identify colonies which were gentamicin sensitive, i.e. the gentamicin cassette flipped out.

To delete END^PaCF1^ or END^PAK^ from CF040 or PAK respectively, the allelic exchange method was used^75^. Briefly, PMQ30 plasmids were first digested with BamHI and HindIII (NEB). Inserts with the required overhangs were ordered as gene fragments from Twist to facilitate cloning. The backbone and inserts were ligated by HiFi DNA Gibson Assembly (NEB), before being transformed into DH5α and selected on gentamicin plates. Plasmids were then extracted using Zyppy miniprep kit (Zymo) as per manufacturer’s protocol and electroporated into SM10 cells. SM10 cells carrying PMQ30 plasmids were then crossed with CF040 or PAK respectively, before selection on VBMM + gentamicin. Gentamicin resistant cassette was then flipped out using FLP recombinase, and negative selection was carried out on no salt LB + 15% sucrose at 30 °C. Colonies were patched onto LB plates with/without gentamicin to identify colonies which were gentamicin sensitive, i.e. the gentamicin cassette flipped out.

### Plate reader assays

Overnight cultures of PAO1 with/without the appropriate p30T constructs were diluted in fresh media with 50 μg/ml gentamicin (if required) at a 1:1000 dilution. Phage lysates were also serially diluted to the appropriate MOI. Diluted cultures (100 μl) were then added to 96-well plates (Thermofisher Scientific) before 10 μl of phage lysates were added and left to grown at 37 °C for 24h in the plate reader (Agilent Technologies), shaking at 425 rpm.

### Plaque assays

Plaque assays were carried out for *P. aeruginosa* and *E. coli*. For *P. aeruginosa*, strains were first grown up overnight with/without antibiotics as required. The overnight cultures (100-150 μl) were then mixed with 0.4% top agar and left to dry for at least 15 minutes. Phage lysates were diluted 10-fold, and 2 μl was spotted onto top agar, before the plates were incubated at 37 °C. No inducers were added to the bottom agar unless as indicated.

For *E. coli*, strains were first grown up overnight with/without antibiotics as required. For experiments with BW25113 carrying p30T constructs, the overnight cultures were then subcultured (1:10) and grown for another 2-3 h, before 100-150 μl were used for mixing with 0.7% top agar and left to dry for at least 15 minutes. For all other experiments with BW25113 and BL21, 100 μl of overnight cultures was mixed with 0.7% top agar. For experiments with DH10B, 0.4% top agar was used to facilitate T4-C plaque formation. Phage lysates were then similarly diluted and spotted before plates were incubated at 37 °C.

### Generating competent *E. coli*

Chemically competent BW25113 and DH10B (from the Bushman lab) was generated as per previous^76^. Briefly, cells were grown to OD_600_ = 0.3 at 37 °C and centrifuged at 4 °C. All remaining steps were done on ice. Cells were washed twice in 50 mM CaCl_2_ with 10 mM Tris pH 7.5 and finally frozen in 50 mM CaCl_2_ with 10 mM Tris pH 7.5 and 15% glycerol with liquid nitrogen before storing at −80 °C. For transformation, cells were subjected to 42 °C heat shock, recovered in LB and plated on 100 μg/ml carbenicillin or 15 μg/ml gentamicin as required.

### Generating recombinant phages

To generate hybrid 14-1 and F8 phages, *P. aeruginosa* PAO1 was first grown overnight, then subcultured and grown to OD_600_ 0.3 at 37 °C, 170 rpm. Phages 14-1 and F8 were then mixed and added to the subculture at MOI = 10. After 10-20 minutes of growth, bacterial cells were centrifuged and resuspended in fresh LB with 10 mM MgSO_4_. After 2 h, a 1% volume of chloroform was added and mixed into the culture. The tubes were then left at 37 °C for 15 min, followed by centrifugation at 10,000 x g, 2 min to remove cell debris. Cells were lysed twice with chloroform, and the supernatant phage lysate was stored at 4 °C.

The hybrid phages generated were then plated onto CF040 carrying Cascade-Cas3 plasmids with guides against 14-1. Large plaques suggestive of hybrid phages rather than escape phages were collected; individual plaques were then propagated in the same CF040-Cas3 background. Phages were then sequenced as per sequencing protocol.

### Generating 14-1 with gp88 frameshifted

Full plate infection using phage 14-1 on PAO1 carrying Cascade-Cas3 plasmid targeting gp88 was carried out in a full plate infection. Individual plaques were then picked and plated on PAO1 with and without END^PaCF1^ to confirm if plaques were true escapers. Phages were then checked for presence of gp88 with primers WX_133/WX_128 and sent for sequencing with Quintara BioSciences.

### Propagation of T4 with different hypermodified/unmodified genomes

Phages T4 wildtype carrying a glucose-hydroxymethyl-cytosine in its genome, T4 carrying hydroxymethyl-cytosine in the genome (T4-HMC) and T4 with unmodified genome (T4-C) were received from the Bushman lab. Strains for phage propagation, CR63 and DH10B, were also received in parallel from the Bushman lab. For wildtype T4 and T4-HMC, phages were propagated on DH10B. For T4-C, phages were first propagated on CR63, before propagating on DH10B to ensure that the resulting phage for experiments have an unmodified genome. All experiments with these three phages were carried out on a DH10B background as per previous^77^.

### Sequencing

CF040 with regions of immune islands deleted were sequenced with Plasmidsaurus with whole bacterial genome sequencing. Sequences were then manually compared to wildtype CF040.

Sequencing of phages was carried out in house as described previously. Briefly, genomic DNA was first extracted using a modified SDS/Proteinase K method. Briefly, 100 μl of phage lysate was mixed with lysis buffer to a final concentration of 10 mM Tris, 1 mM EDTA, 0.5% SDS before incubating at 37 °C for 30 min, followed by 55 °C for 30 min – 1 h. DNA was then purified using DNA Clean and Concentrator kit and quantified with Nanodrop. 20-100 ng genomic DNA was used to prepare WGS libraries using Illumina DNA Prep Kit. Subsequently, PCR indexing-amplification of tagmented DNA was performed using 2x Phusion Master Mix (NEB) and custom-ordered indexing primers, amplified for 12 cycles. Libraries were further purified by agarose gel electrophoresis and purified with Zymoclean Gel DNA Recovery Kit (Zymo) as per manufacturer’s instructions, Trimmed reads were assembled de-novo with SPADES^78^. All hybrid phage genomes were then queried for 14-1/F8 SNPs in the genome using the command-line ‘grep’ function.

### qPCR

PAO1 with/without END^PaCF1^ were grown overnight in LB + 10 mM MgSO_4_. Cultures were then subcultured at a 1:50 dilution and grown to OD_600_ = 0.4. F8 were then added to cultures at MOI of approx. 1. Samples (500 μl) were then taken at t = 0, 15, 30 and 60 min post infection and washed once with cold 100 μl PBS (Gibco). Samples were then frozen on dry ice, before genomic DNA was extracted using the modified SDS/Proteinase K method as previously described. Genomic DNA (2 – 10 ng) was then used for qPCR with Luna Universal qPCR Master mix (NEB), with *rpoD* primers and primers WX_Q4F/WX_Q4R, on the CFX Connect Real-Time PCR Detection System (Bio-Rad) at UCSF.

### Restriction digest

100 ng of F8/14-1 gDNA were incubated with 1 μl of restriction enzyme (NEB) as stated in 50 μl rCutSmart reaction at 37 °C, 1h. For BmrI, NEB buffer r2.1 was used. Reactions were then run on 0.8% agarose gel in TAE buffer.

### Protein purification

BL21 carrying plasmids were first grown overnight at 37 °C in LB 150 μg/ml carbenicillin. Cells were then subcultured into 100 ml LB 150 μg/ml carbenicillin and grown to log phase for 1.5-2 h. Inducer (1mM IPTG) was then added and the cultures were left to grow at 18 °C overnight.

The next day, cultures were centrifuged at 4000 x g for 10 min. Cells were then resuspended in lysis buffer, before four rounds of sonication. Lysates were then put through a nickel column and washed four times, each with 4 volumes of resin. Proteins were then eluted with elution buffer three times, each with an equivalent volume of resin. The lysates, wash 4 and elution 1 were collected. SDS-PAGE gels were then run using Tris-Glycine SDS running buffer (Bio-Rad) at 170 V for 45 min and stained with Direct Blue. Gels were then imaged with the Gel Doc.

### Characterization of phage-modified nucleosides

LC-MS/MS fragmentation analysis for identifying the chemical identity of the base modification in F8 and 14-1 genomes were carried out as per previous^48^. Briefly, F8 and 14-1 DNA were digested to nucleosides overnight with Nucleoside Digestion Mix in 1x Nucleoside Digestion Mix Reaction buffer (NEB). The resulting DNA nucleosides were then analyzed using Agilent 6490 Triple quadrupole LC/MS system. The chromatography was performed using a Waters XSelect HSS T3 C18 column (2.1 × 100 mm, 2.5 μm particle size) operated at a flow rate of 0.6 mL/min with a linear gradient of aqueous buffer (10 mM aqueous ammonium formate, pH 4.4) and methanol over 6 minutes. MS/MS fragmentation spectra were obtained by collision-induced dissociation in the positive product ion mode with the following parameters: gas temperature 200 °C, gas flow 14 L/min, nebulizer 45 psi, sheath gas temperature 350 °C, sheath gas flow 11 L/min, capillary voltage 2 kV, nozzle voltage 1.5 kV, fragmentor voltage 380 V and collision energy 5–65 V.

### Bioinformatics analysis

Amino acid alignments were done using ClustalOmega^79^ and webLOGO was used to identify the consensus sequence. To determine genomic neighborhoods, amino acid sequence of END^PaCF1^ were input into webFLAGS^38^. For determining the CTD domain, the amino acid sequence was input into HHpred^58^. For identifying potential genes involved in synthesizing hypermodified genomes, Domainator^50^ and Foldseek^51^ were used. Alignment of phage genome sequences from NCBI was done with Clinker^80^.

To determine total number of homologs, PSI-Blast was carried out as previously described^2^. Briefly, homologs of END^PaCF1^ were first identified using PSI-Blast with the maximum number of target sequence set at 5,000 and E-value cut-off of 0.005. Two more iterations were then carried out, and the retrieved sequences were aligned using MAFFT^81^. The resulting alignment was then visualized using Jalview^82^, and entries which resulted in large gaps in the alignments were discarded. Due to the large difference in size, entries larger and smaller than 500 amino acid sequences were further differentiated.

For analysis of other homologs with shared domain, two iterations of PSI-BLAST was ran against complete genomes. The best 760 hits were analyzed (E-value ≤ 10^−12^). Only hits with domains aligning to either the N or the C terminal domain of END^PaCF1^ were selected. The iENDOIII CTD domain was further ran against the clustered NR and domain organizations of retrieved proteins with significant sequence similarity were examined.

Protein structures were predicted using AlphaFold3^59^, and visualized using PyMoL^83^.

## Author contributions statement

W-X. Y., T. A. K., A. W., A. E. G. generated strains, conducted experiments and analyses. Y-J. Lee conducted nucleoside analysis. K. S. M. conducted bioinformatic analysis of homologs with shared domains. W-X. Y. drafted the initial manuscript, and T. A. K., B. C., K. S. M., E. V. K., P. R. W. and J. B. D. reviewed and edited the text. J. B. D. and P. R. W. provided overall supervision and secured funding.

## Acknowledgements

We would like to thank Sukrit Silas and Horia Todor for providing *E. coli* strain BW25113 and the Basel phages, the Brouns lab at TU Delft for the vB_PaeM_FBPa21 phage (annotated in this paper as vB-Pa21) and the Bushman lab at University of Pennsylvania for T4 phages with different genome hypermodifications.

---

**Extended Data Figure 1:**
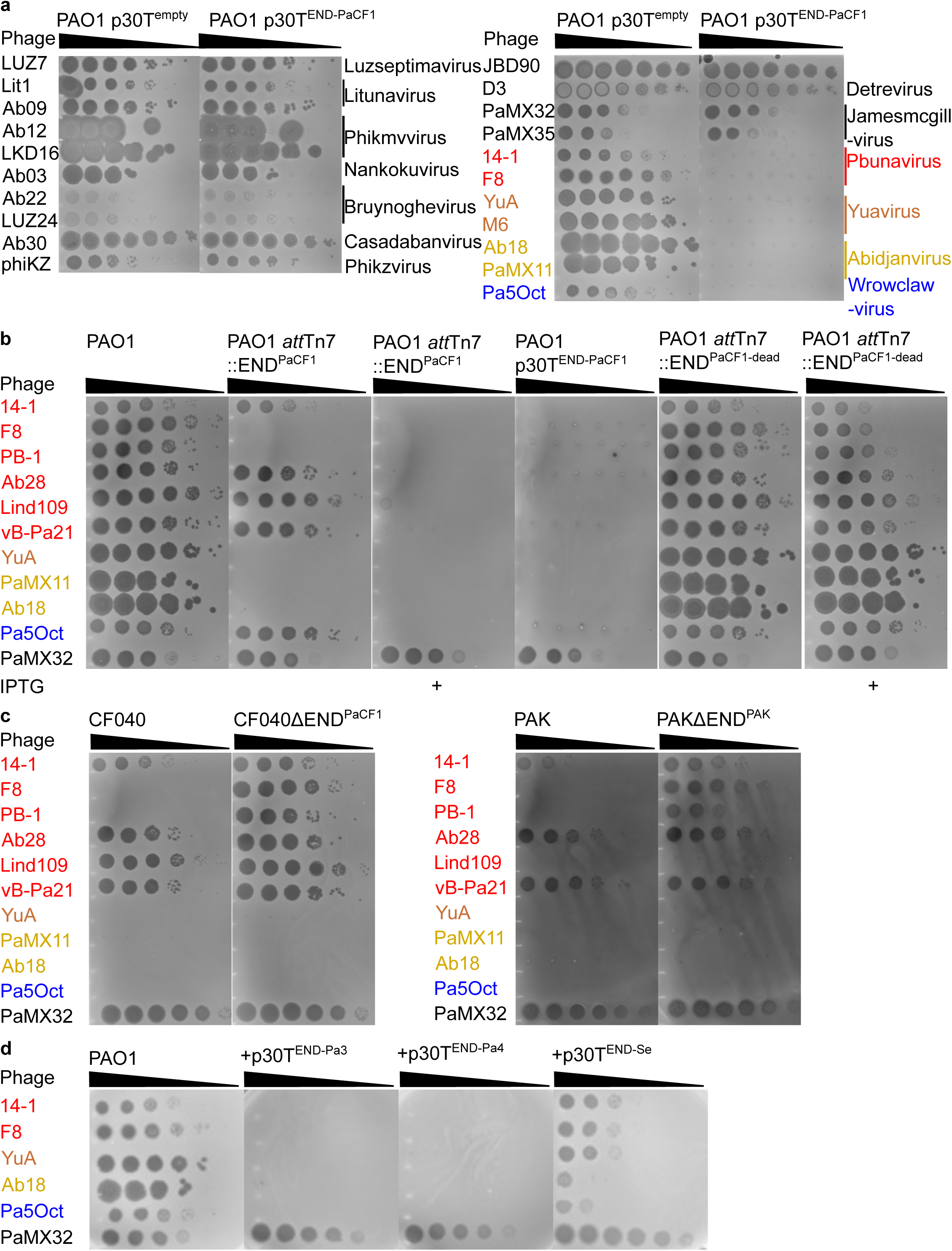
END^PaCF1^ and other homologs targets F8-like and YuA-like phages, and is functional in native strains CF040 and PAK. **a**, Phages across different phage families were plaqued onto PAO1 carrying p30T^END-PaCF1^. Empty p30T plasmid was included as a control; p30T plasmid was used to maximise expression of END^PaCF1^. **b**, Plaque assays for all tested phages with increasing induction of END^PaCF1^. 1mM IPTG was used to induce *att*Tn7::END^PaCF1^. Plaque assays are representative of three repeats. **c**, Phages from F8 family, YuA family and Pa5Oct were plaqued onto CF040 and PAK with and without END^PaCF1^/ END^PAK^ deleted. **d**, Plaque assays for PAO1 carrying p30T^END-PaCF1^ homologs. PaMX32 was included as a phage control. Plaque assays shown are representative of at least three repeats.

**Extended Data Figure 2:**
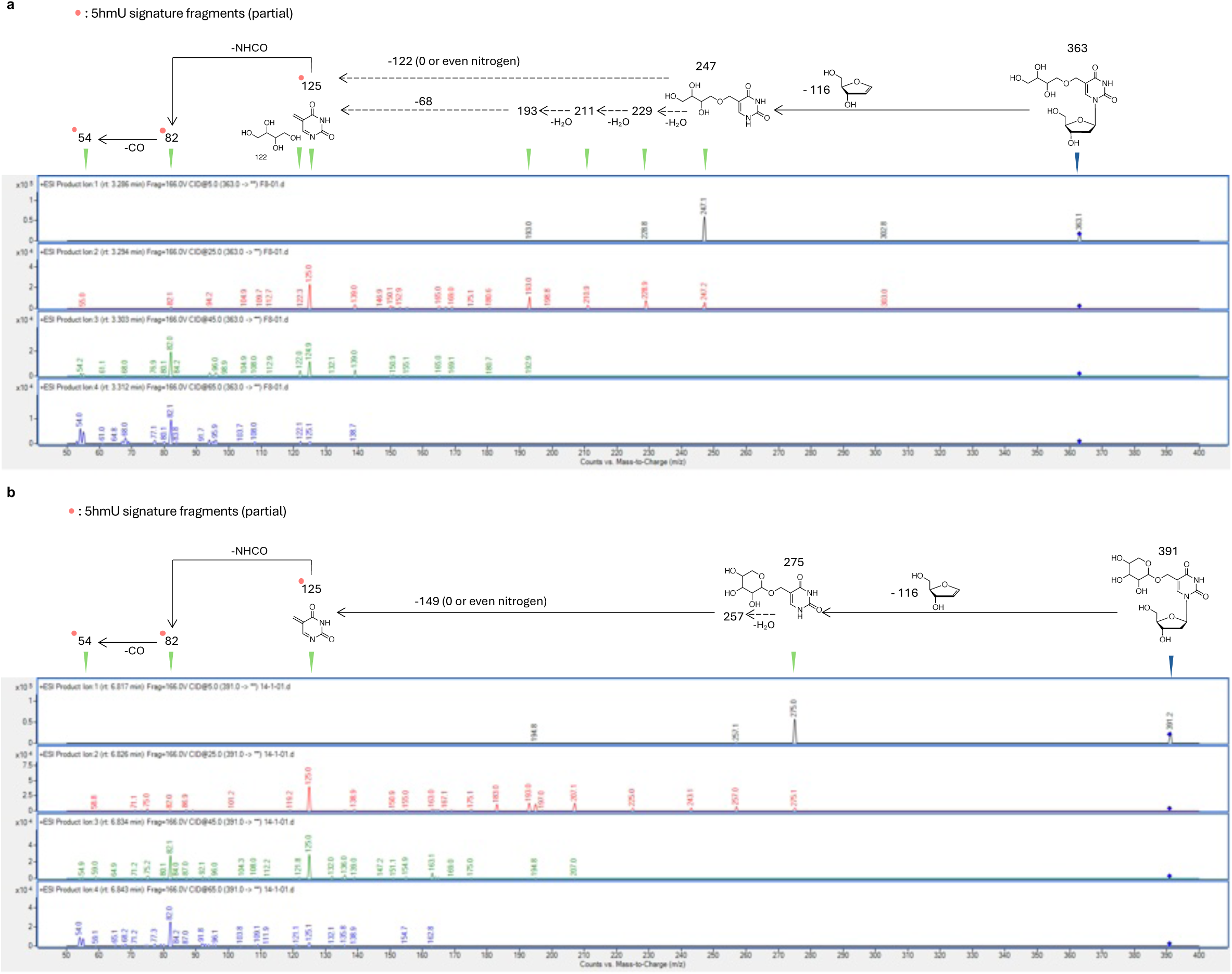
+ESI-MS/MS fragmentation spectra of nucleoside from phages F8 and 14-1. Product-ion spectra with collision-induced dissociation (CID) energies of 5, 25, 45 and 65 is as shown for **a,** F8 and **b,** 14.1

**Extended Data Figure 3:**
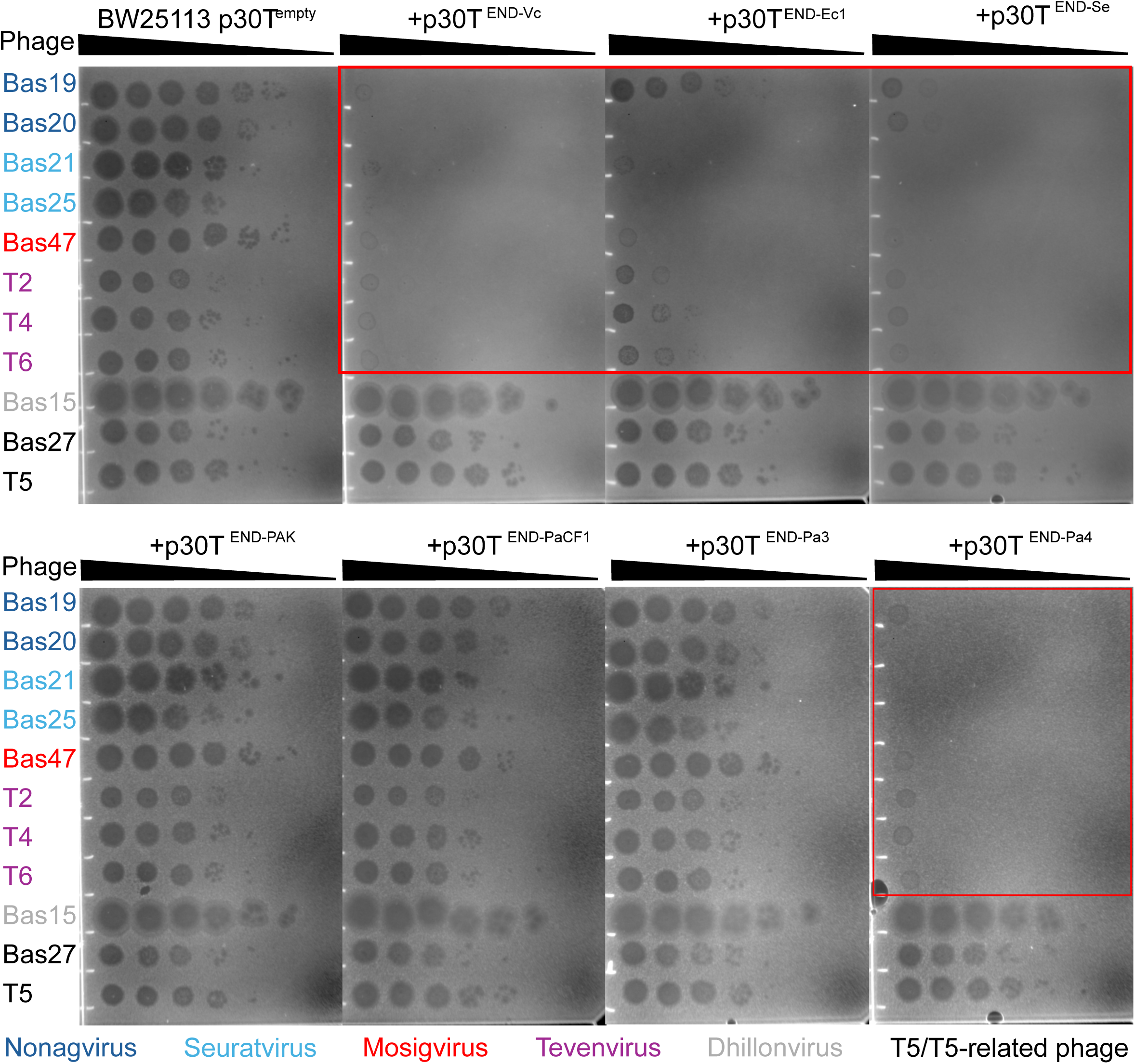
Homologs of END^PaCF1^ inhibit *E. coli* phages with hypermodified genomes. Panel of *E. coli* phages from the Basel collection was tested against END^PaCF1^ homologs across different bacteria. Red boxes highlight phages which showed a decrease in titer in the presence of the defense system. T5 and T5 related phages, which are known to have non-modified genomes, are used as controls.

**Extended Data Figure 4:**
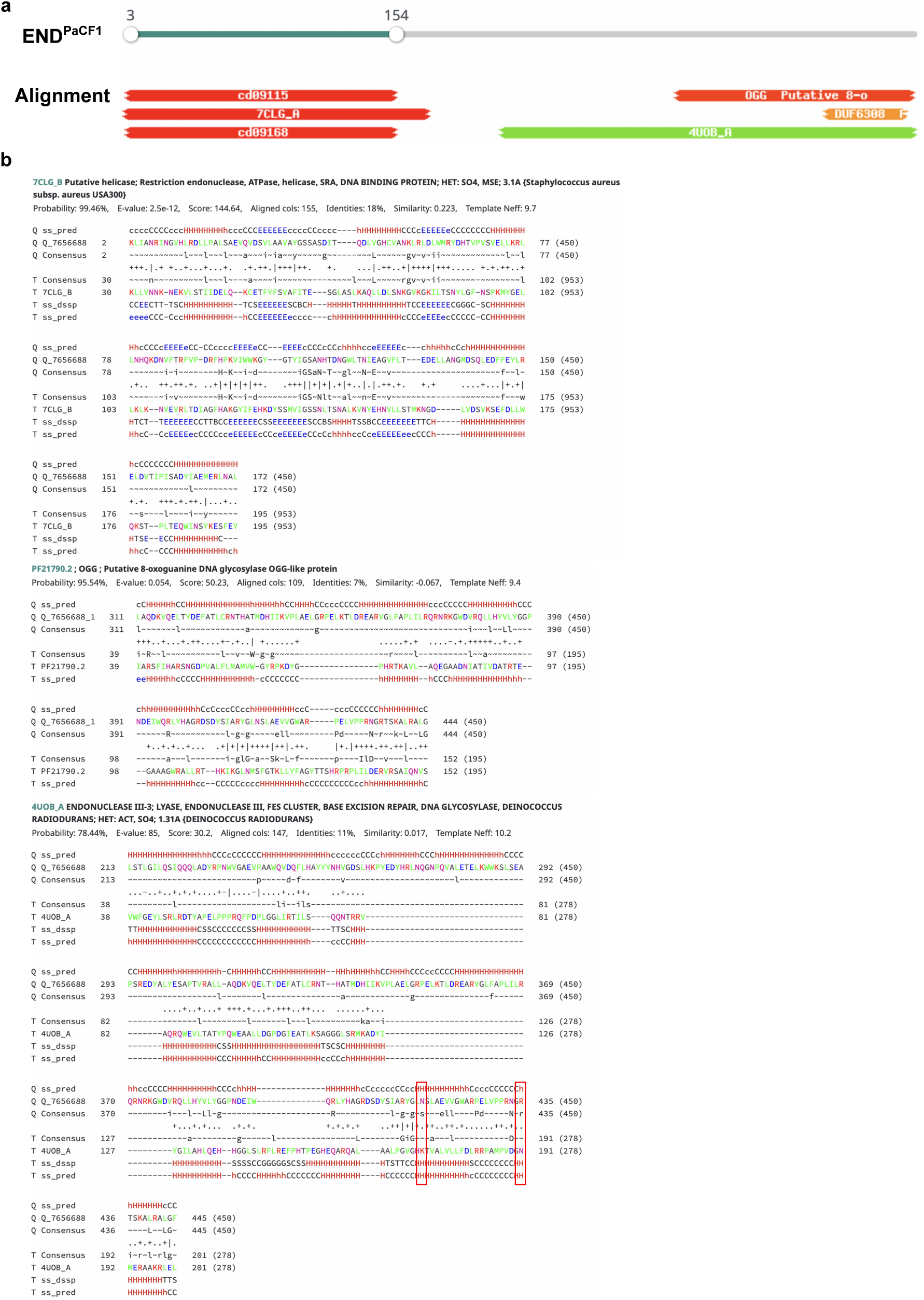
HHpred domain analysis of END^PaCF1^. **a**, Alignment output from HHpred. Only the top 3 alignment of each domain is shown. **b**, Details of alignment of END^PaCF1^ with characterized proteins. First alignment is for END^PaCF1^ NTD and PLD nuclease domain from ATP-dependent restriction endonuclease SauUSI (PDB: 7CLG_A). Second alignment is for END^PaCF1^ CTD and 8-oxo-guanine DNA glycosylase OGG-like (PF21790). Third alignment is for END^PaCF1^ CTD and endonuclease III (PDB: 4UOB). For the third alignment, the catalytic residues of 4UOB is shown boxed in red.

**Extended Data Figure 5:**
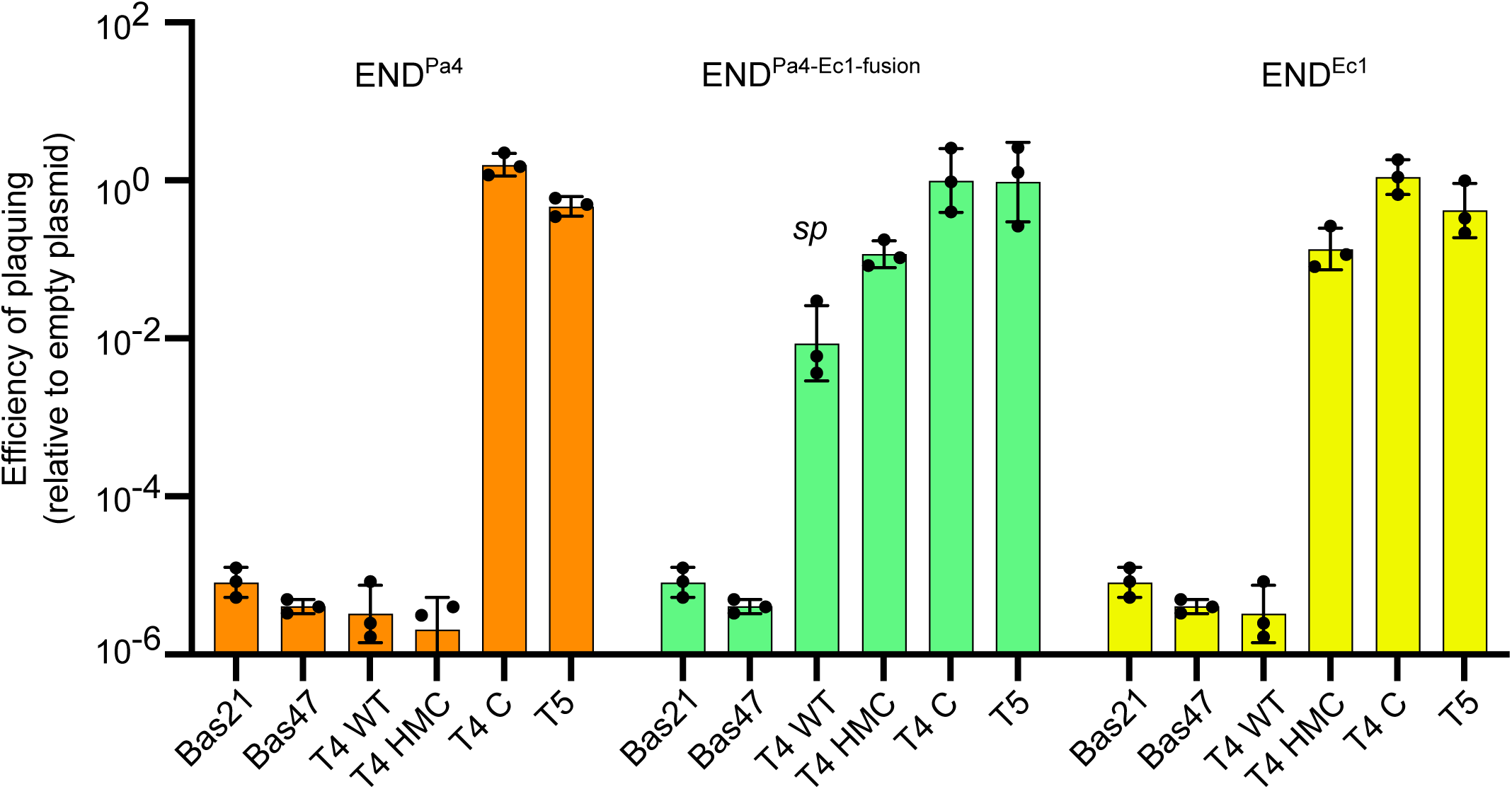
Histogram of plaque assays in *E. coli* DH10B with END^Pa4^, END^Ec1^ and END^Pa4−Ec1−fusion^. Panel of *E. coli* phages were tested against different END nuclease constructs. T5 is included as a negative control. HMC = hydroxymethylcytosine, C= unmodified cytosine. sp stands for small plaques (see Figure 4e).

**Extended Data Figure 6:**
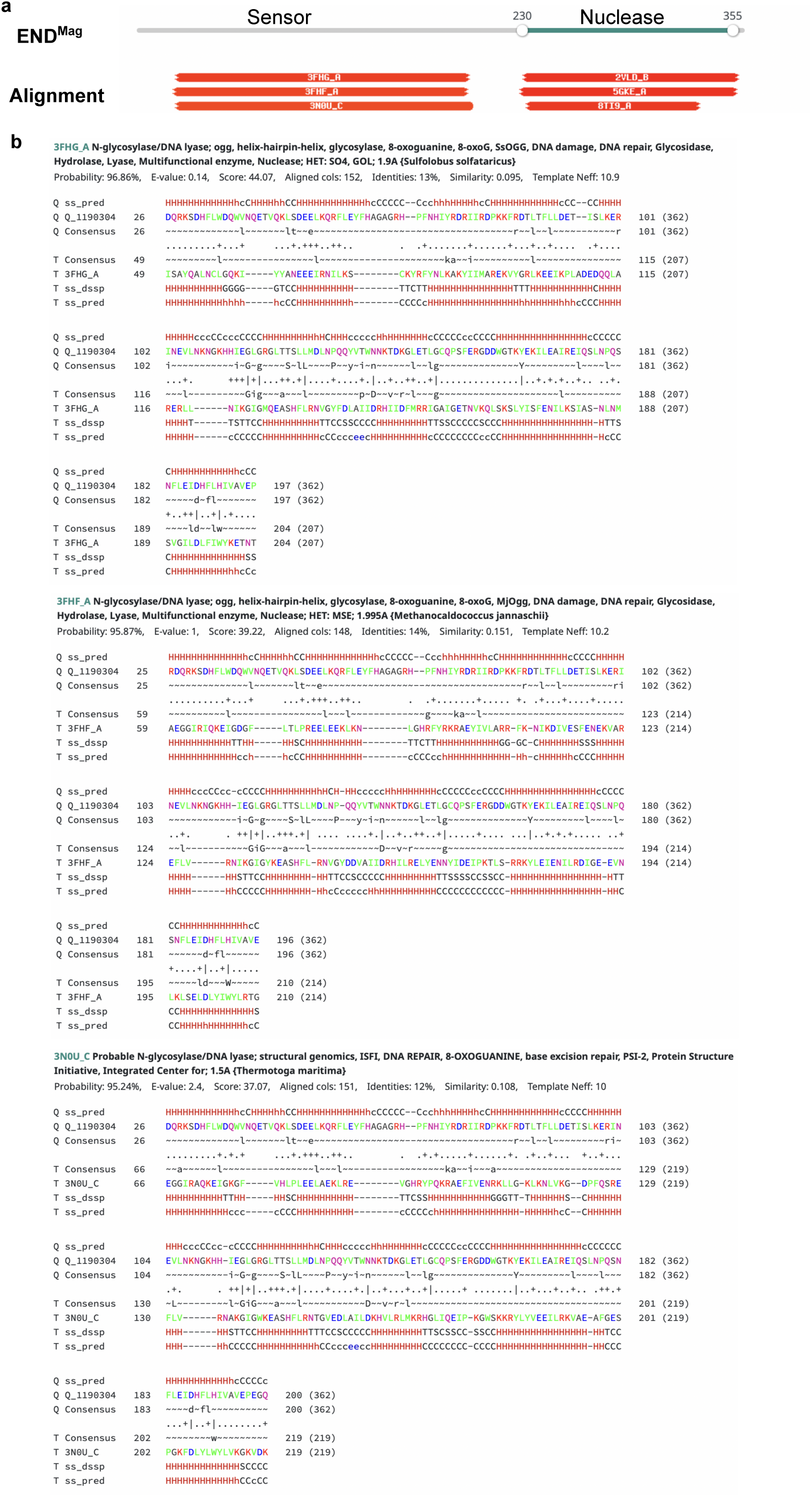
HHpred domain analysis of END^Mag^. **a**, Alignment output from HHpred. Only the top 3 alignment of each domain is shown. **b**, Details of alignment of END^Mag^ sensor domains with characterized proteins. First alignment is for END^Mag^ with *Sulfolobus solfataricus* Ogg1 (PDB: 3FHG). Second alignment is for END^Mag^ with *Methanocaldococcus jannashii* Ogg1 (PDB: 3FH3). Third alignment is for END^Mag^ with *Thermotoga maritima* Ogg1

**Extended Data Figure 7:**
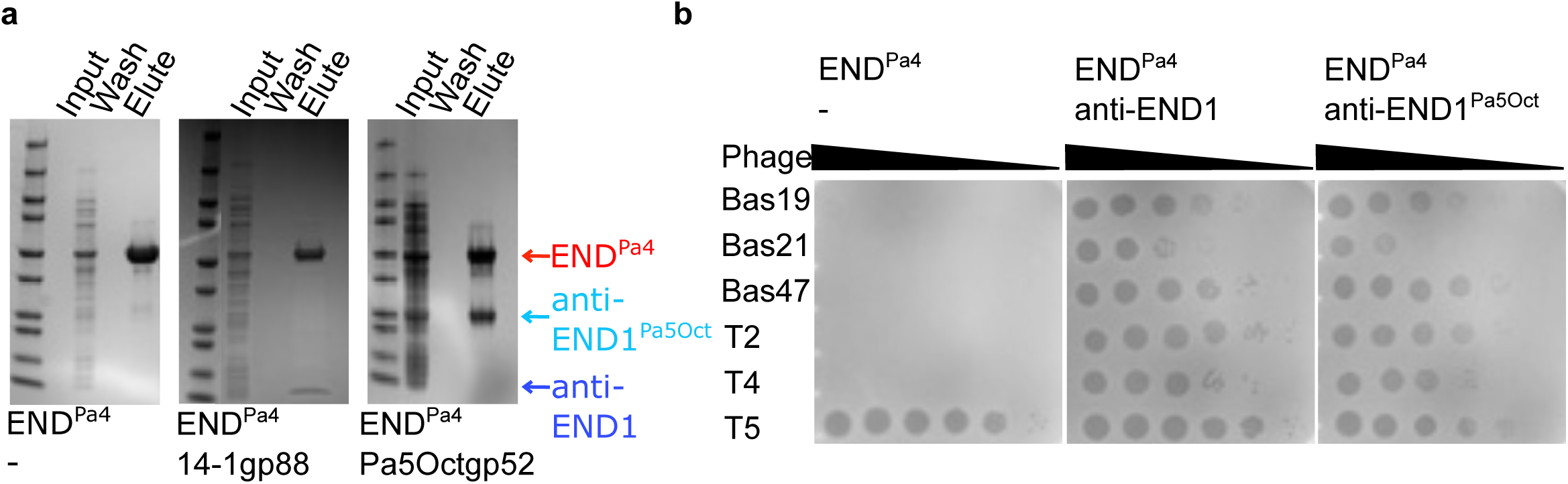
END^Pa4^, a distant homolog of END^PaCF1^, also binds to anti-END inhibitors. **a,** Pulldown of inhibitors in the presence of defense system. Only END^Pa4^ is His-tagged; controls are shown in Figure 5. **b,** Plaque assay of different hypermodified phages in *E. coli*, in the presence of END^Pa4^ with/without anti-END inhibitors.

**Extended Data Table 1:** Putative roles of defene-associated genes and other genes indicated in Figure 1b.

| Notation | Coordinates | strand | DefenseFinder | HHpred |
| --- | --- | --- | --- | --- |
| gene 1 | 6354280..6356735 | - | Type IV RM (note no nuclease) |  |
| gene 2 | 6356736..6358225 | - | Mokosh |  |
| gene 3 | 6358210..6361595 | - | Mokosh (note no nuclease) |  |
| gene 4 | 6361602..6363005 | - |  | S/T kinase |
| gene 5 | 6363011..6363703 | - |  | Protein serine/threonine phosphatase |
| gene 6 | 6363703..6364374 | - |  | Von Willebrand factor type A (vWA) domain |
| gene 7 | 6365945..6367771 | - | OLD/Gabija |  |
| gene 8 | 6376092..6377573 | - |  | RtcR, Regulator of RNA terminal phosphate cyclase |
| gene 9 | 6376092..6377573 | - |  | RtcR, Regulator of RNA terminal phosphate cyclase |
| gene 10 | 6379523..6380257 | - |  | N-terminal glycosyltransferase |
| gene 11 | 6380320..6381444 | - |  | ComF-like phosphoribosyltransferase |
| gene 12 | 6381447..6382313 | - |  | Histidinol phosphatase (PHP family) |

**Extended Data Table 2:**
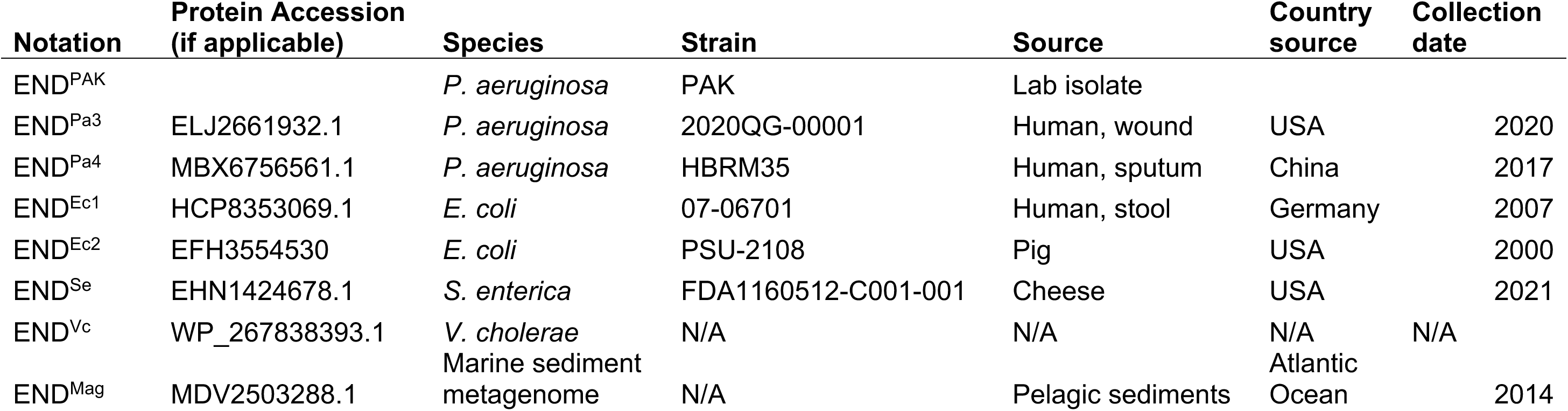
END nucleases tested experimentally in the paper.

| Notation | Protein Accession<br>(if applicable) | Species | Strain | Source | Country<br>source | Collection<br>date |
| --- | --- | --- | --- | --- | --- | --- |
| END<sup>PAK</sup> |  | *P. aeruginosa* | PAK | Lab isolate |  |  |
| END<sup>Pa3</sup> | ELJ2661932.1 | *P. aeruginosa* | 2020QG-00001 | Human, wound | USA | 2020 |
| END<sup>Pa4</sup> | MBX6756561.1 | *P. aeruginosa* | HBRM35 | Human, sputum | China | 2017 |
| END<sup>Ec1</sup> | HCP8353069.1 | *E. coli* | 07-06701 | Human, stool | Germany | 2007 |
| END<sup>Ec2</sup> | EFH3554530 | *E. coli* | PSU-2108 | Pig | USA | 2000 |
| END<sup>Se</sup> | EHN1424678.1 | *S. enterica* | FDA1160512-C001-001 | Cheese | USA | 2021 |
| END<sup>Vc</sup> | WP_267838393.1 | *V. cholerae* | N/A | N/A | N/A | N/A |
| END<sup>Mag</sup> | MDV2503288.1 | Marine sediment<br>metagenome | N/A | Pelagic sediments | Atlantic<br>Ocean | 2014 |

**Extended Data Table 3:** List of restriction enzymes and the corresponding recognition sequence as indicated on NEB website.

| **Restriction enzyme** | **Recognition sequence** |
| --- | --- |
| BsgI | GTGCAG(16/14) |
| EcoRI | G/AATTC |
| FspI | TGC/GCA |
| HindIII | A/AGCTT |
| BmrI | ACTGGG(5/4) |
| ApaLI | G/TGCAC |
| NdeI | CA/TATG |
| XbaI | T/CTAGA |
| DraI | CACNNN/GTG |
| BsaI | GGTCTC(1/5) |

**Supplementary Data Table 1:**
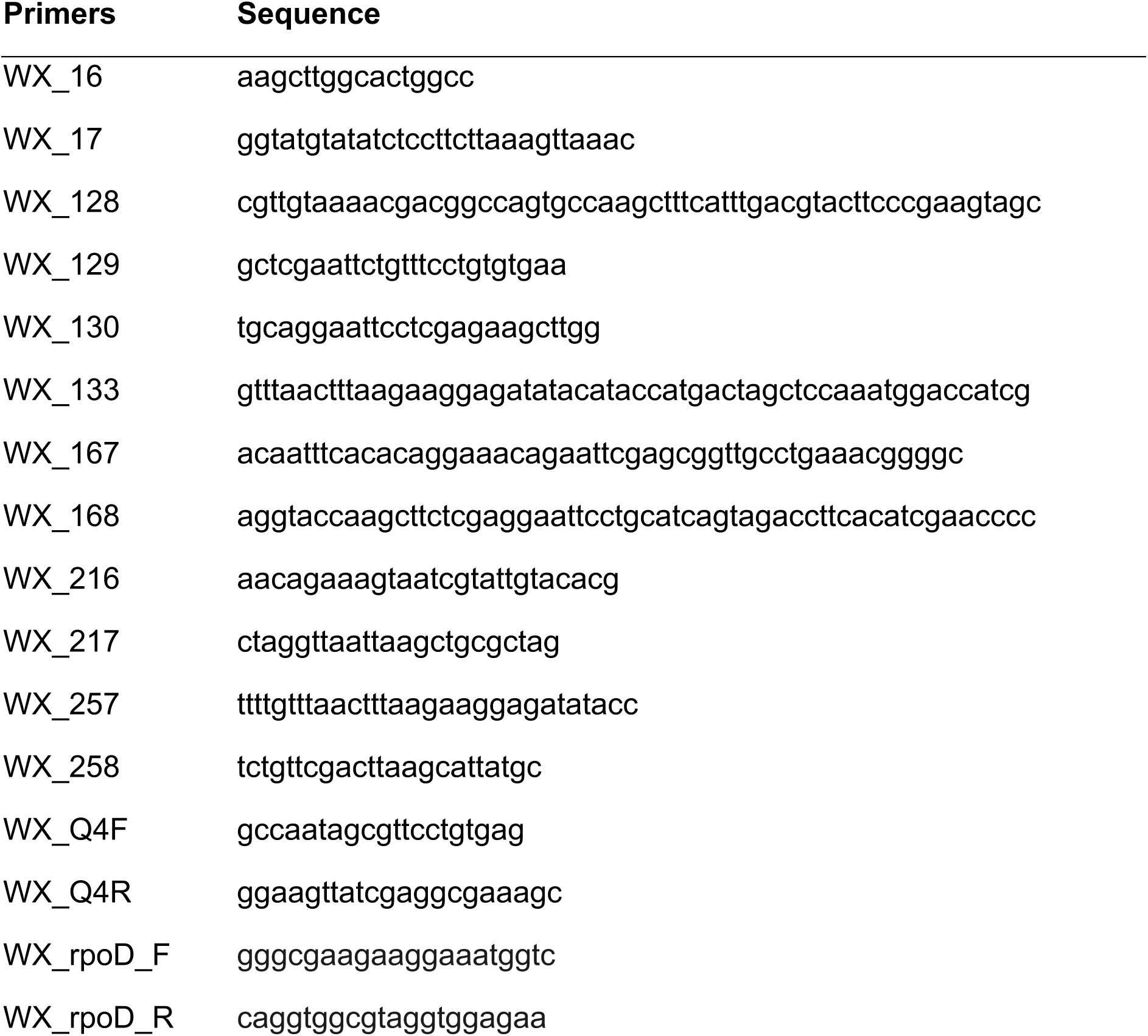
List of primers used.

